# Genome Architecture Leads a Bifurcation in Cell Identity

**DOI:** 10.1101/151555

**Authors:** Sijia Liu, Haiming Chen, Scott Ronquist, Laura Seaman, Nicholas Ceglia, Lindsey A. Muir, Walter Meixner, Pin-Yu Chen, Gerald Higgins, Pierre Baldi, Steve Smale, Alfred Hero, Indika Rajapakse

## Abstract

Genome architecture is important in transcriptional regulation and study of its features is a critical part of fully understanding cell identity. Altering cell identity is possible through overexpression of transcription factors (TFs); for example, fibroblasts can be reprogrammed into muscle cells by introducing MYOD1. How TFs dynamically orchestrate genome architecture and transcription as a cell adopts a new identity during reprogramming is not well understood. Here we show that MYOD1-mediated reprogramming of human fibroblasts into the myogenic lineage undergoes a critical transition, which we refer to as a bifurcation point, where cell identity definitively changes. By integrating knowledge of genome-wide dynamical architecture and transcription, we found significant chromatin reorganization prior to transcriptional changes that marked activation of the myogenic program. We also found that the local architectural and transcriptional dynamics of endogenous MYOD1 and MYOG reflected the global genomic bifurcation event. These TFs additionally participate in entrainment of biological rhythms. Understanding the system-level genome dynamics underlying a cell fate decision is a step toward devising more sophisticated reprogramming strategies that could be used in cell therapies.

## INTRODUCTION

A comprehensive understanding of cell identity, how it is maintained and how it can be manipulated, remains elusive. Global analysis of the dynamical interplay between genome architecture (form) and transcription (function) brings us closer to this understanding (Rajapakse and Groudine, 2011). This dynamical interaction creates a genomic signature that we can refer to as the four-dimensional organization of the nucleus, or 4D Nucleome (4DN) (Chen et al., 2015; Dixon et al., 2015; Fortin and Hansen, 2015; Krijger et al., 2016). Genome technologies such as genome-wide chromosome conformation capture (Hi-C) are yielding ever higher resolution data that give a more complete picture of the 4DN, allowing us to refine cell types, lineage differentiation, and pathological contributions of cells in different diseases. High temporal resolution on a global scale provides key insight into biological processes. Understanding the dynamical process of cellular reprogramming is of interest in regenerative medicine, as it can improve our ability to generate cells for repair and regeneration of tissue in injury and disease.

Pioneering work by Weintraub et al. showed that reprogramming fibroblasts into muscle cells was possible through overexpression of a single transcription factor (TF), MYOD1, thereby demonstrating that a different cell identity could supersede an established one (Weintraub, 1993; Weintraub et al., 1989). In 2007, when Yamanaka and colleagues reprogrammed human fibroblasts into an embryonic stem cell-like state with four TFs, POU5F1 (OCT4), SOX2, KLF4, and MYC, they showed that a pluripotent state could also supersede an established cell identity (Takahashi et al., 2007). These remarkable findings demonstrate the possibilities of controlling the genome and the cell identity through TFs. However, how TFs dynamically orchestrate genome architecture and transcription as a cell changes identity during reprogramming is not fully understood.

One exciting finding in recent reports is that Hi-C contact maps can be used to divide the genome into two major compartments, termed A and B (Chen et al., 2015; Lieberman-Aiden et al., 2009). Compartment A is associated with open chromatin (transcriptionally active), and compartment B with closed chromatin (transcriptionally inactive). It was shown that during differentiation, structurally defined regions of active/inactive gene expression (i.e. A/B compartments) changed to facilitate the different expression of a new cell state (Chen et al., 2015; Dixon et al., 2015, 2012; Lieberman-Aiden et al., 2009).

Previously we introduced a new technique from spectral graph theory to partition the genome into A/B compartments and identify topologically associating domains (TADs) (Chen et al., 2016). This motivated us to study the 4DN from a network point of view, where nodes of the network correspond to genomic loci that can be partitioned at different scales such as 1-Mb region, 100-kb region, gene level, and TAD level. The edges of the network indicate contact between two loci, with contact weights given by Hi-C entries. From the network perspective, A/B compartments are identified as distinct connected components of a network. As will be demonstrated here, other properties of the network topology, such as network centrality measures, can be extracted from Hi-C data to yield further information about chromatin spatial organization.

The utility of network centrality allows one to identify nodes (e.g., genomic loci) that play influential topological roles in the network (Newman, 2010). A number of centrality measures exist, each specialized to a particular type of nodal influence. For example, degree centrality characterizes the local connectedness of a node as measured by the number of edges connecting to this node, while betweenness centrality is a global connectedness measure that quantifies the number of times a node acts as a bridge along the shortest path between two other nodes. Eigenvector centrality is a neighborhood connectedness property in which a node has high centrality if many of its neighbors also have high centrality. Google’s PageRank algorithm uses a variant of eigenvector centrality (Lohmann et al., 2010). Our work builds biological connections between Hi-C contact maps and network centrality.

In this paper we investigated dynamics of topological features of genome architecture and explored how they varied with transcription during MYOD1-mediated reprogramming of human fibroblasts into the myogenic lineage. Sampling across a time course during reprogramming, we captured architecture by Hi-C, transcription by RNA-seq, and protein content by proteomics. By combining different centrality measures we found important Hi-C features largely overlooked in previous studies, and this approach facilitated coordinated form-function analysis of chromatin conformation and gene expression in genome-wide data. The concept of bifurcation is introduced to describe a critical transition from one cell identity to another. Analyses of form-function dynamics revealed a bifurcation in space-time 32 hours after activation of exogenous MYOD1 in fibroblasts that suggested a definitive transition into the myogenic lineage. During the reprogramming process, we found that chromatin reorganization occurs prior to changes in transcription. Moreover, we found robust synchronization of circadian gene expression, and determined that these genes are downstream targets of MYOD1, suggesting MYOD1 feedback onto circadian gene circuits. After the bifurcation, MYOG was associated with synchronization of a subset of important myogenic TFs. These findings support roles for MYOD1 and MYOG in entraining biological rhythms. Finally, our analysis of genomic regulatory elements such as chromatin remodeling genes, super enhancer regions and microRNAs provides additional clues toward understanding system-wide dynamics during reprogramming.

## RESULTS

### MYOD1-mediated Direct Reprogramming: Revisiting Weintraub

We converted primary human fibroblasts into the myogenic lineage using the TF and master regulator MYOD1, similar to Weintraub’s studies on myogenic reprogramming (Weintraub, 1993) (Figure 1A). In the system, MYOD1 initiates the transcriptional program that turns muscle cell precursors into multinuclear muscle fibers. Fibroblasts were transduced with a lentiviral construct that expressed human MYOD1 fused with the tamoxifen-inducible mouse ER(T) domain (L-MYOD1) (Kimura et al., 2008). With 4-hydroxytamoxifen (4-OHT) treatment, transduced cells showed nuclear translocation of L-MYOD1 and morphological changes consistent with myogenic differentiation (Figure S1). We then validated the activation of two key myogenic genes downstream of *MYOD1* (*MYOG* and *MYH1*) (Figure 1B). These results demonstrate successful conversion of fibroblasts into the myogenic lineage by L-MYOD1. Subsequent analyses were carried out on 4-OHT treated, transduced cells, sampling at 8-hour (hr) intervals for RNA-seq, small RNA-seq, and Hi-C analyses, and at 24-hr intervals for proteomics (Figure 1C). We focused on 12 samples (-48, 0, …, 80 hrs), capturing genome architecture (form) through Hi-C and genome-wide transcription (function) through RNA-seq. The resulting time series data on chromatin conformation and gene expression can be studied at different scales (Figure 1D).

**Figure 1:**
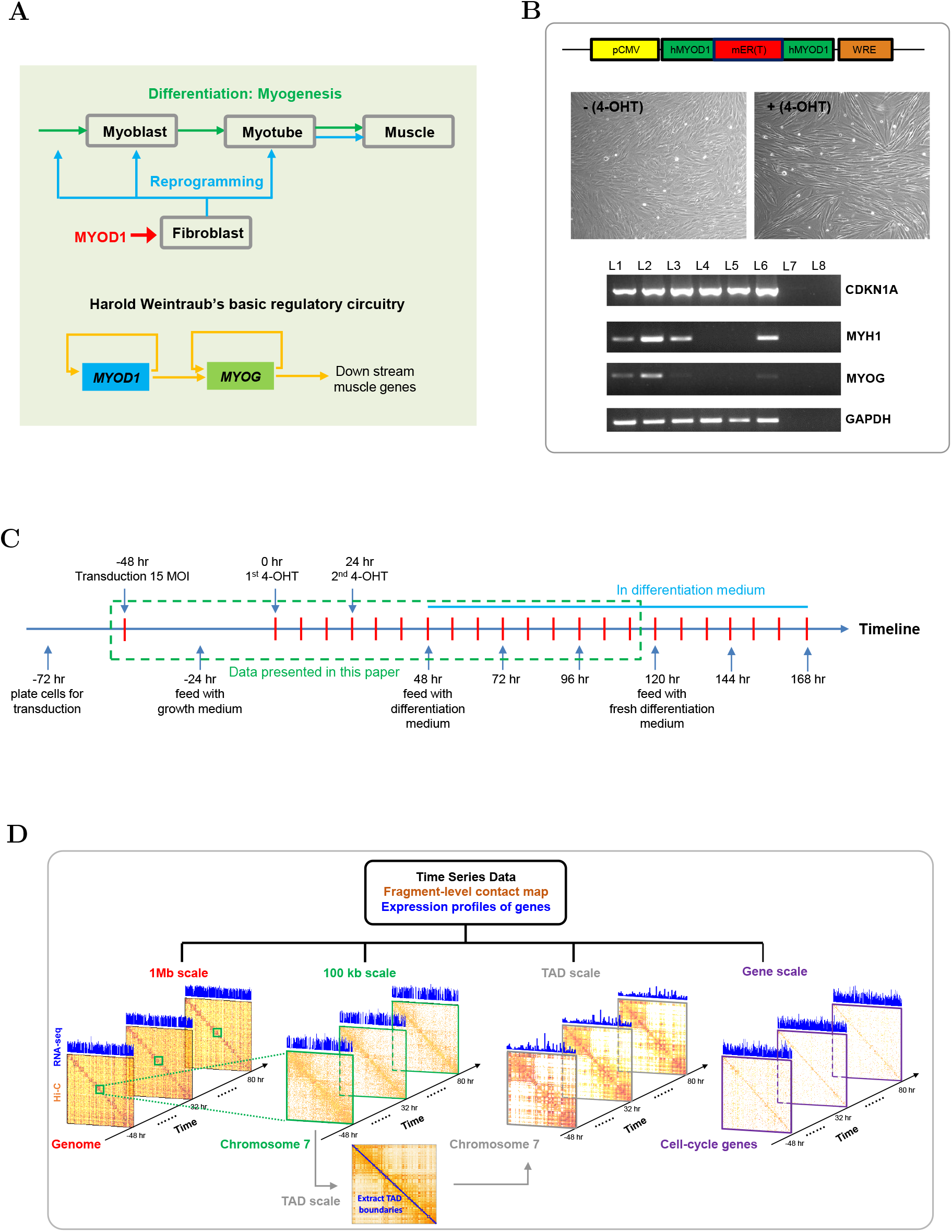
Experimental design to explore cellular reprogramming with time-evolving genome architecture (form) and gene expression (function). (A) *Top*: Potential cell state transition pathways for MYOD1-mediated fibroblast to muscle cell reprogramming. *Bottom*: Basic gene regulatory circuitry for myogenesis. (B) *Top*: The cassette for myogenic reprogramming lenti-construct, expressing a fusion protein with the mouse mER(T) domain (red box) inserted within the human MYOD1 (green boxes) between amino acids 174 and 175. *Middle*: Light microscope images of cells without (left) or with (right) 4-OHT treatment at differentiation day 3. *Bottom*: RT-PCR validation of gene expression at day 3. Lanes L1/2/3/6 are samples transduced with L-MYOD1, no transduction (L4) or lenti-vector only (L5), L7 is RT-negative control, and L8 is water for PCR negative control. Two key MYOD1 downstream genes, *MYOG* & *MYH1* are activated by the expression of L-MYOD1. *GAPDH* is used as internal control, and *CDKN1A* (P21) is universally expressed. (C) Time course of MYOD1-mediated reprogramming. The time window outlined in green corresponds to time points at which both genome architecture and transcription are captured by Hi-C and RNA-seq. (D) Scale-adaptive Hi-C matrices and gene expression. The considered scales include 1 Mb, 100 kb, TAD and gene level.

### A Bifurcation Delineates Emergence of a New Cell Identity

We first explored whether our data would reveal any critical state transitions as cells adopted a new identity. For this analysis, we interpreted Hi-C and RNA-seq data as measurements of dynamic networks, where Hi-C contact maps depict network topologies, and RNA-seq data characterize the function of nodes (Figure 2A). With the aid of network representation, we captured multiple topological properties of genome architecture using the concept of network centrality (STAR Methods).

**Figure 2:**
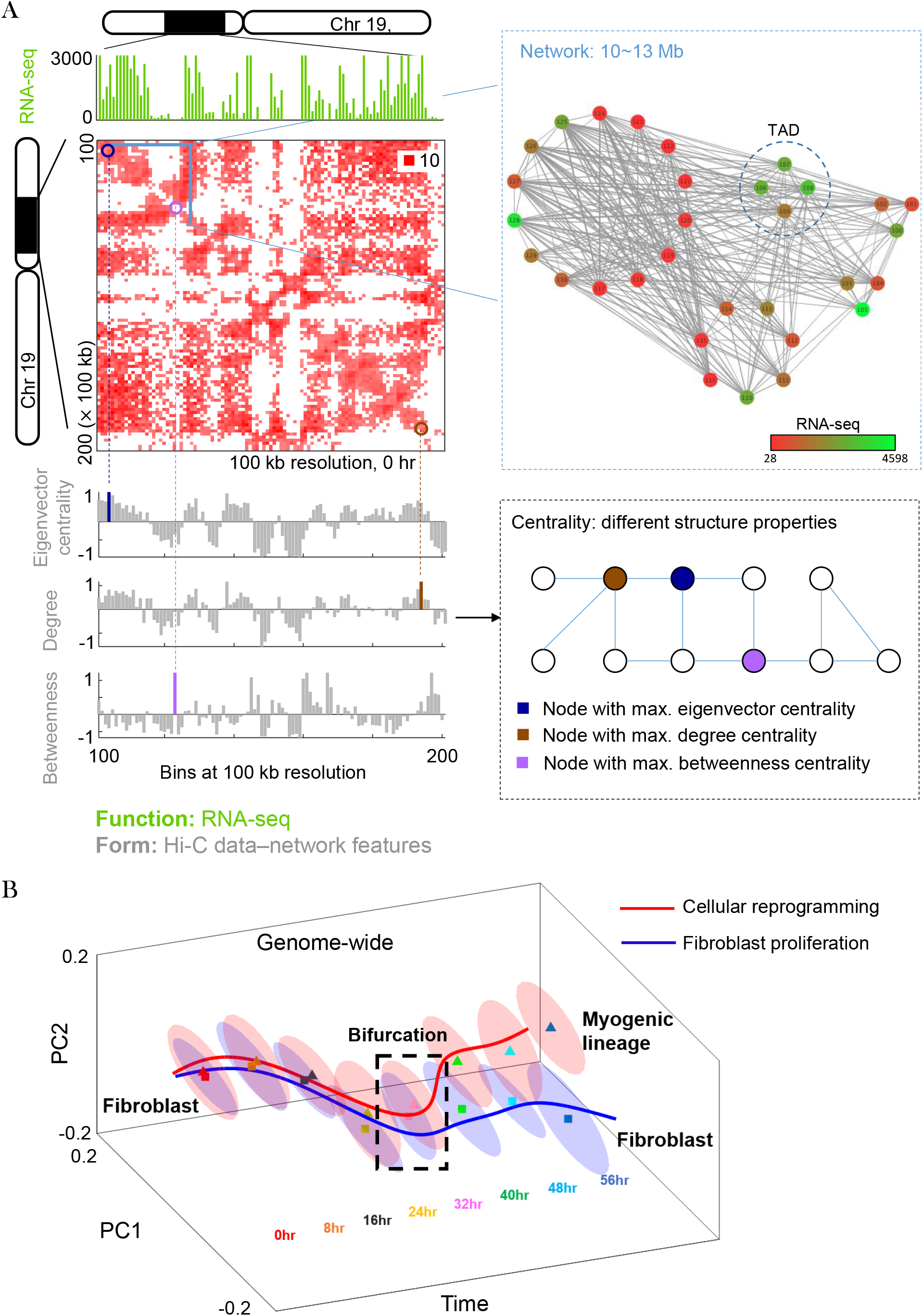
Network representation of form-function time series data and bifurcation detection. (A) Mapping genomic form (Hi-C contact map) and function (RNA-seq) to network structure and node dynamics. *Top left*: Hi-C contact map (Toeplitz normalized, STAR Methods) and RNA-seq at 100-kb resolution for chromosome 19. *Top right*: Network representation in which edge width indicates the Hi-C contact number and node color implies the magnitude of RNA-seq FPKM value. *Bottom left*: Network features given by eigenvector centrality, degree centrality and betweenness centrality scores. The bars marked by different colors correspond to maximum centrality values. *Bottom right*: An illustrative network under different centrality measures. (B) Cell state trajectory of MYOD1-mediated reprogramming and fibroblast proliferation (Chen et al., 2015). Ellipsoids represent low-dimensional data representations obtained by applying Laplacian eigenmaps (STAR Methods) to network form-function features. The branching trajectory shows a bifurcation point at 32 hrs with P-value < 0.01.

We found that network centrality such as eigenvector centrality, degree centrality and betweenness centrality quantified the importance of genomic loci from different angles, each of which provides a signature of genome structure (Figures 2A and 3). Specifically, eigenvector centrality identified structurally defined regions of active/inactive gene expression, namely, A/B compartments. Additionally, eigenvector centrality yielded a higher correlation with transcriptional activity than conventionally defined A/B compartments, which are derived from the first principal component (PC1) of a spatial correlation Hi-C matrix (Figures 3A and S2) (Lieberman-Aiden et al., 2009). Betweenness centrality recognized regions that switched A/B compartment assignment between time steps. The values of betweenness at A/B switched regions were significantly higher than other centrality measures (Figure 3B), and 70% of switched regions were located at boundaries between open and closed chromatin compartments (Figure S3). This finding suggests that the boundary regions, which connect A/B compartments, serve as bridge-nodes [what is a bridge-node, and can you interpret what that means in the genome? Could you say that high betweenness is a way to identify the boundaries, which may tell you something about the biology?] in the network, and therefore correspond to high betweenness values. Furthermore, we identified 546 genes (Table S1) from A/B switched loci between 0 and 40 hrs which had at least two-fold difference in expression levels. There were 395 genes with increased expression levels and 151 genes with decreased expression. By performing functional annotation for these genes (Huang et al., 2009), we identified 36 genes enriched (FDR < 0.005) under UniProtKB keywords ‘cell cycle’ (Bairoch et al., 2005). Among these genes, 22 are involved in cell division, and 19 are related to mitosis. This suggests that in our reprogramming regime, the cells are proliferating while incubating in growth medium (before induction of differentiation).

**Figure 3:**
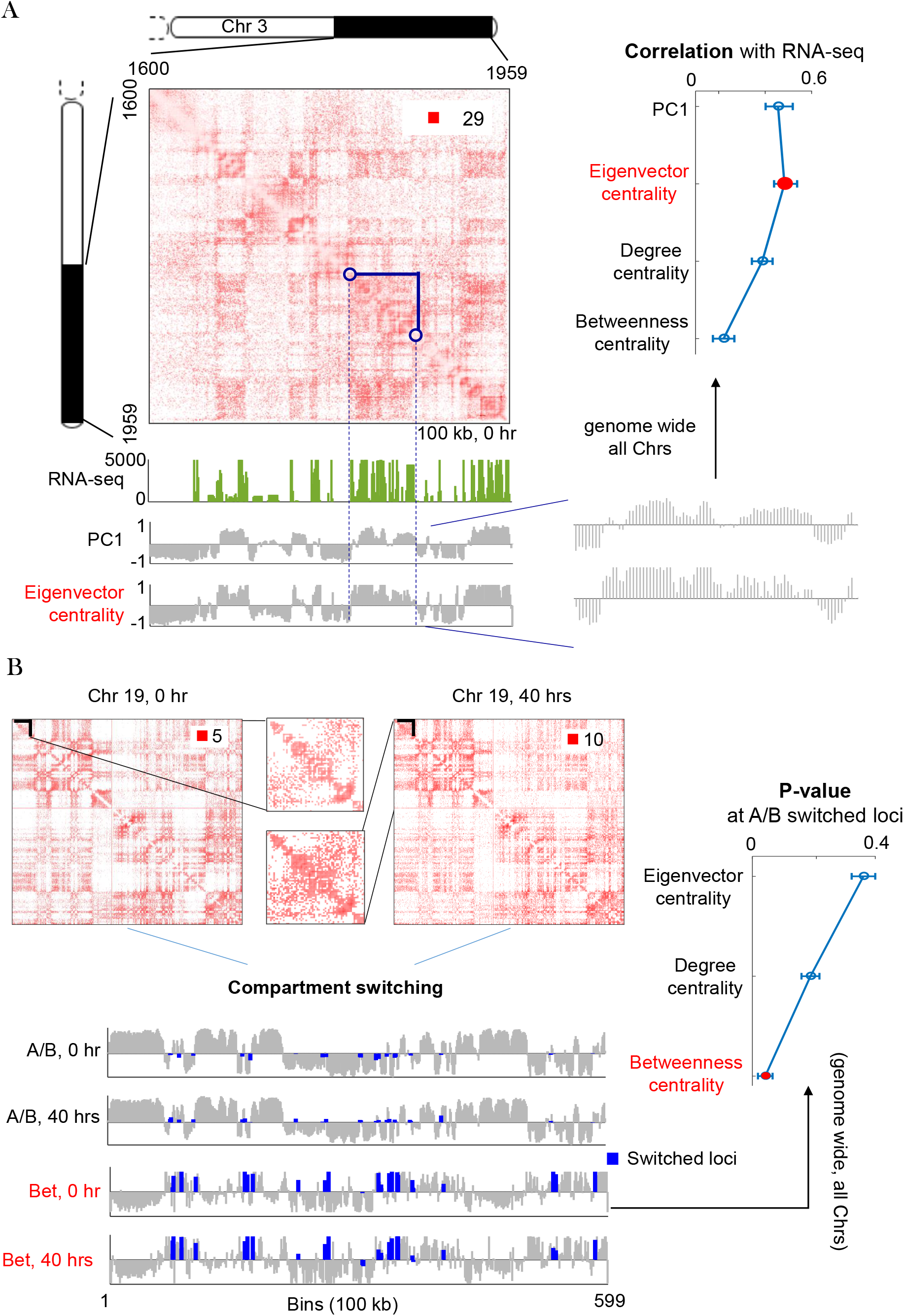
Biological interpretation of network centrality measures. (A) Eigenvector centrality indicates chromatin compartments, termed A and B. *Top left*: Hi-C contact map of chromosome 3 at 100 kb resolution. *Bottom left*: RNA-seq, the first principal component (PC1) of Hi-C spatial correlation matrix, and eigenvector centrality (in terms of its z-score). *Right*: Correlation between RNA-seq and PC1 and centrality features extracted from Hi-C data for all chromosomes. Eigenvector centrality is a better indicator for chromatin compartments. (B) Betweenness centrality indicates A/B switched loci. *Top left*: Hi-C contact map of chromosome 19 (100 kb resolution) at time instants 0 and 40 hrs. *Bottom left*: A/B partition and betweenness centrality (in terms of its z-score) at 0 and 40 hrs. The blue color represents A/B switched bins from 0 to 40 hrs. The switched loci tend to have large betweenness centrality scores. *Right*: Significance of betweenness centrality at A/B switched loci. The P value is determined by comparing the average betweenness value at A/B switched bins to a random background distribution of other centrality values under the same number of bins.

After integrating centrality-based network features with the function of nodes (namely, gene expression), we extracted a low-dimensional representation of genome-wide form-function data using the dimension reduction technique of Laplacian eigenmaps (STAR Methods). The resulting form-function representation, fitted by a minimum volume ellipsoid (MVE) (STAR Methods), showed distinct configurations at different time points (Figure 2B). We compared our reprogramming dataset to proliferating fibroblast data obtained using similar methods over a 56-hr time course (Chen et al., 2015). In the latter experiment, the cells were cell cycle and circadian rhythm synchronized before introduction to full growth medium, with time point collection of RNA-seq and Hi-C every 8 hours. We discovered a divergence of these two datasets at 32 hrs with *P*=0.0048 (STAR Methods), suggesting a cell state transition at this time that marked an abrupt shift in the genomic system from its prior state (fibroblast) to a new state (myogenic). The P value is determined by a Hotelling’s T-squared test associated with the null hypothesis that the centroids of MVEs under the reprogramming and proliferation data are identical at a given time point. Our result aligns with a branching trajectory that can be represented using the concept of *bifurcation* (Borchert and Slade, 1981): there exists a bifurcation point in the sense that two simultaneous trends starting from the same state (fibroblast) become separated from each other, toward two stable equilibria (fibroblast and myogenic).

To further examine the pathway into the myogenic lineage, we focused on topologically associating domains (TADs) (Dixon et al., 2012) containing genes related to myogenesis (myoblast, myotube, and skeletal muscle; Table S2). Our data suggest transit directly to a more differentiated myotube-like state without a myoblast-like stage (Figure S4). Bypass of an intermediate stage of differentiation during reprogramming is also supported by expression patterns of key myogenic genes: *MYF5*, *MYOD1*, and *MYOG* (Weintraub et al., 1991). It is known from natural myogenic differentiation (Bentzinger et al., 2012) that *MYF5* is expressed first in myoblasts, while *MYOD1* and *MYOG* are up-regulated in the more differentiated myotubes. During reprogramming, *MYF5* expression was not detectable across time points, whereas *MYOD1* and *MYOG* expression increased after the bifurcation at 32 hrs.

Next, we found that the bifurcation event was reflected in the local dynamics of *MYOD1* and *MYOG*. As noted before, endogenous *MYOD1* and *MYOG* expression were first detected around the bifurcation point. In addition, the bifurcation was identified in Hi-C contact maps of *MYOD1* and *MYOG* (Figure S5). Here the difference between gene-level Hi-C matrices at successive time points revealed a pattern strikingly similar to what was found in genome-wide dynamics: a critical transition was observed at the bifurcation that delineates emergence of a new cell identity.

Taken together, we interpret three phases for reprogramming from our data: fibroblast, bifurcation, and myogenic. A phase transition between the bifurcation and the myogenic phase can be detected from analysis of genome-wide dynamics or local dynamics of key regulators, *MYOD1* and *MYOG*.

### Form Precedes Function

Given the cell state trajectory, it was unclear whether MYOD1-mediated reprogramming induced rewiring of genome architecture prior to the role of MYOD1 in mediating muscle gene transcription, or vice versa (Kosak and Groudine, 2004; Rajapakse and Groudine, 2011). To answer this question, we focused on form and function dynamics of 22083 genes genome-wide, where the form is depicted by inter-gene contact maps (STAR Methods), and the function corresponds to RNA-seq FPKM values (Figure 4A). The form-function evolution is then evaluated by temporal differences of network centrality features (extracted from inter-gene contact maps) and gene expression, named as form/function temporal difference score (TDS; STAR Methods). Based on TDS at successive time points (Figure 4B), we found that a significant form change at 8 hrs preceded a significant function change at 16 hrs. For deep understanding of form-function evolution during the reprogramming process, we applied K-means clustering (with two clusters) on both form and function data. This was done to identify subsets of genes that yielded the most significant temporal change (Figure 4C and D). From this analysis we found that genes contained within each cluster of high TDS, which are responsible for function and form change, at most have 20% overlap (Table S3). This suggests that the mechanism of form evolution could be different from that of function evolution, and that these two mechanisms are steered by different sets of genes. Furthermore, we investigated five gene modules extracted from Gene Ontology (GO): fibroblast, myotube, skeletal muscle, cell cycle, and circadian genes (Table S2). By contrasting our reprogramming dataset with data on human fibroblast proliferation (Chen et al., 2015), we found that the pattern of form-function evolution during reprogramming is quite different from fibroblast proliferation (Figure 4E). Consistent with findings represented in Figure 4B, the effects of nuclear reorganization were detectable prior to transcription changes, that is, form preceded function. Our results support the speculation that [I think ‘‘suggest” is too strong here, because you don’t demonstrate that these changes are required for the transcriptional changes to occur. If you don’t like the word speculate, you could use “we propose” or similar] that chromatin architectural changes facilitate the orchestrated activation of transcriptional networks associated with adoption of a new cell identity.

**Figure 4:**
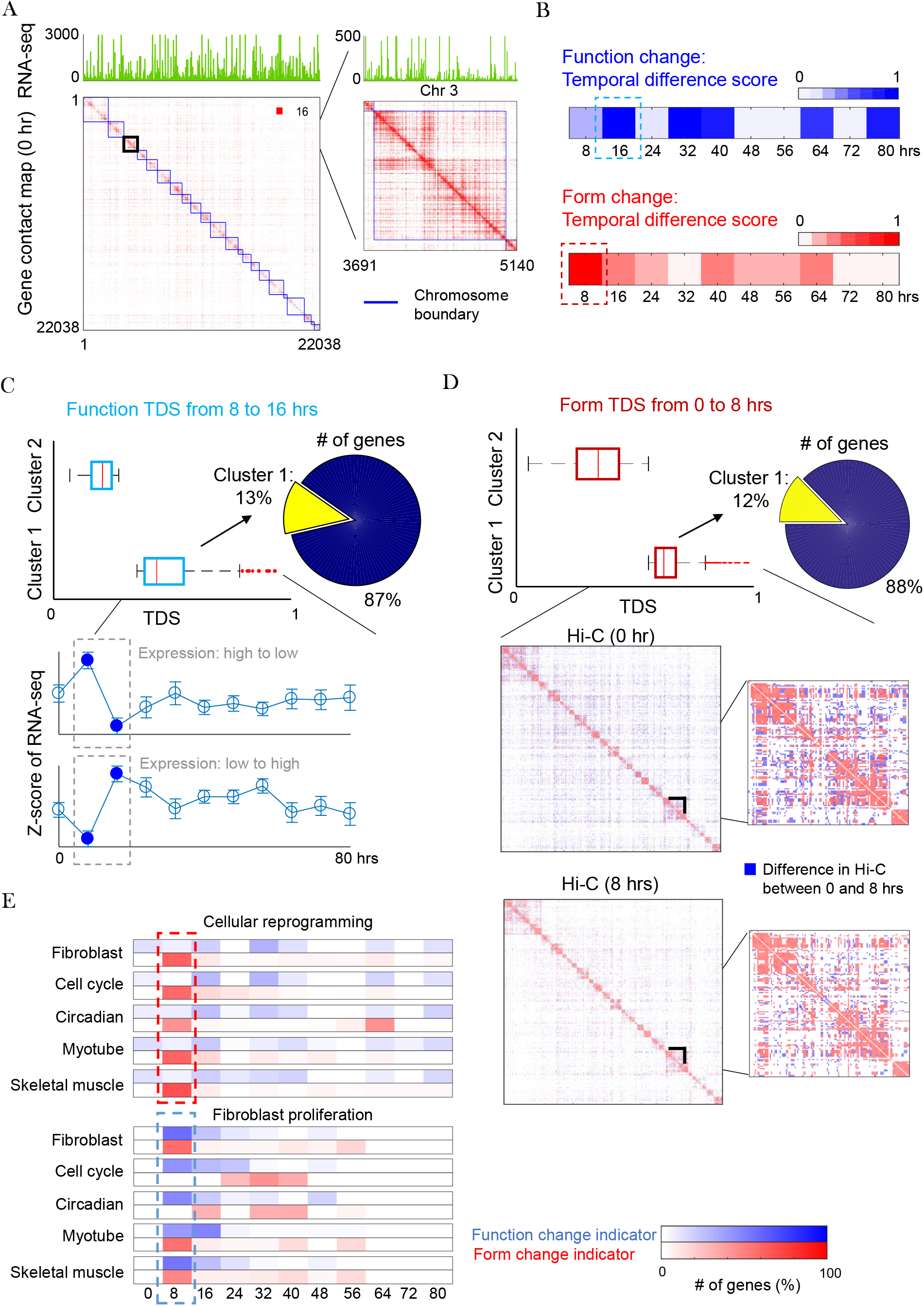
Genome-wide dynamical form-function relationship. (A) Genomic form and function given by Hi-C contact map and RNA-seq. (B) Function and form change at successive time points evaluated by temporal difference score (TDS; STAR Methods) of RNA-seq and network centrality features of Hi-C data, respectively. The significant form change (at 8 hrs) occurs prior to the function change (at 16 hrs). (C) Illustration of function TDS from 8 to 16 hrs. Genes are divided into two clusters by applying K-means to their TDS values. Cluster 1 contains genes with the largest temporal change in RNA-seq. The gene expression can either decrease or increase from 0 to 16 hrs. (D) Illustration of form TDS from 0 to 8 hrs. Two gene clusters are obtained by applying K-means to their TDS values. Hi-C contact maps associated with a subset of genes in cluster 1 are shown from 0 to 8 hrs, where the blue color indicates the Hi-C difference between two time points. (E) Form-function change indicators for gene modules of interest during cellular reprogramming (top) and fibroblast proliferation (bottom), respectively. The number (%) of genes with significant form-function change is differentiated by color.

Given a gene module (focusing on fibroblast or muscle related genes), we found that genes that are associated with significant form change at 8 hrs and function change at 32-40 hrs were in the majority (> 30%) of each of gene modules, while only a small portion of these genes (< 5%) had the same significant form-function change during human fibroblast proliferation (Figure S6A). We extracted the genes (77 in fibroblast module and 72 in muscle module; Table S4) that were only responsible for significant change during cellular reprogramming and those that were less active during fibroblast proliferation. This gives a set of core genes shown by its distinct form-function evolution during reprogramming (Figure S6B, C and D). The statistical significance on the temporal change of the identified genes was evaluated as P < 0.05 by comparison to fibroblast proliferation data (STAR Methods).

### Portrait of 4DN: Proliferation versus Reprogramming

We next introduce the concept of a ‘portrait of 4DN’, which provides a simple quantitative assessment of form-function dynamical relationships at the chromosome level. Specifically, on a 2D plane, we designate one axis as a measure of form in terms of network connectivity (STAR Methods), and the other as a measure of function in terms of the average RNA-seq FPKM value. The portrait of 4DN is then described by a form-function domain, made up of 8 time points [0,56] (hrs) for each chromosome (Figure 5A). Although there exists similar patterns in the portrait of 4DN between the cellular reprogramming and the human fibroblast proliferation datasets, a difference between the bifurcation point (32 hr) of the two settings was observed (Figure 5B). Also, the centroid of the fitted form-function ellipsoid (MVE estimate; STAR Methods) for each chromosome shifts between cellular reprogramming and fibroblast proliferation. Such a shift can be decomposed into the horizontal change and the vertical change, where the former describes the form change, and the latter corresponds to the function change. We found that most chromosomes undergo more form change (86.4%) compared to function change (13.8%) (Figure 5C). Furthermore, the area of the chromatin ellipse is able to characterize the variance (uncertainty) of 4DN (Figure 5C). We observed that ellipsoids associated with cellular reprogramming had larger volumes than those under human fibroblast proliferation. This implies that cellular reprogramming leads to a more complex dynamical behavior, indicated by a space-time bifurcation and phase transition during this process (Figure 2D).

**Figure 5:**
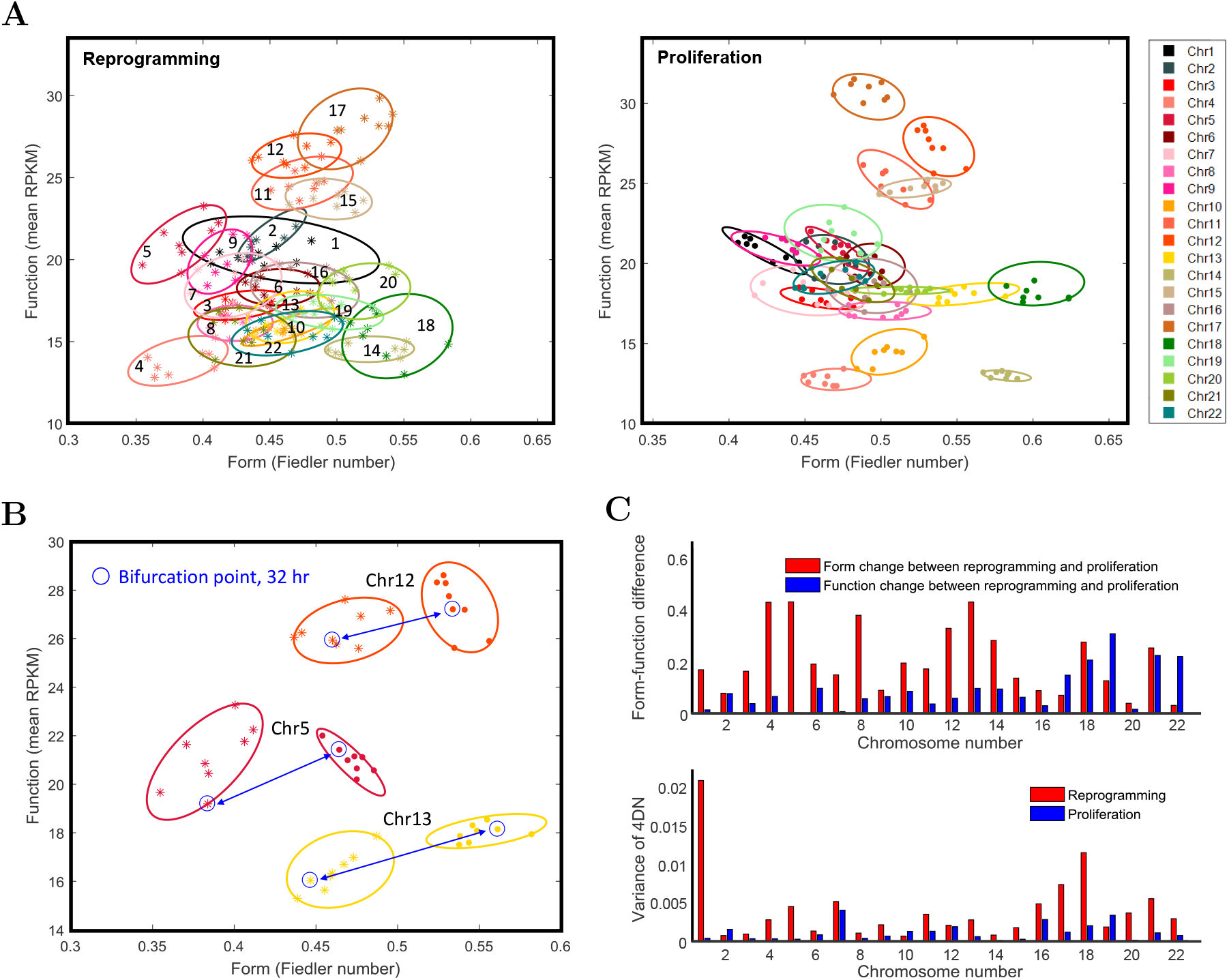
Portrait of 4DN under cellular reprogramming and human fibroblast proliferation. (A) Portrait of 4DN in the context of reprogramming and proliferation, respectively. It is described by a form-function domain (2D), made up of 8 time points, for each chromosome. The fitted ellipsoid is obtained from MVE estimate (STAR Methods). (B) Shift of form-function domains of chromosomes at the bifurcation point (32 hr). Chromosomes 5, 12 and 13 yield the top three most significant changes. (C) Differences between cellular reprogramming and fibroblast proliferation, indicated by centroids and volumes of form-function ellipsoids for each chromosome. *Left*: Comparison between form change (horizontal shift) and function change (vertical shift) for each chromosome. *Right*: Variance of 4DN, given by volumes of chromosome ellipsoids under different cell dynamics.

### MYOD1-mediated Synchronization of Circadian Rhythms

A number of studies have explored the link between MYOD1 and circadian genes ARNTL and CLOCK, revealing that ARNTL and CLOCK bind to the core enhancer of the *MYOD1* promoter and subsequently induce rhythmic expression of *MYOD1* (Andrews et al., 2010; Zhang et al., 2012). Here we discovered that upon *MYOD1* activation, circadian genes exhibited robust synchronization in gene expression, suggesting MYOD1 feedback onto the circadian gene network. Further inspection showed that core circadian genes (Table S2) that contain E-boxes displayed the most profound synchronization initially, starting with an uptick in gene expression just after MYOD1 activation (Figure 6A-D). JTK_CYCLE (Hughes et al., 2010) confirmed our observation; all E-box circadian genes were found to have a synchronized period of 24 hrs, with a maximum lag of 4 hrs between any genes, with the exception of *CRY1* (Table S5).

**Figure 6:**
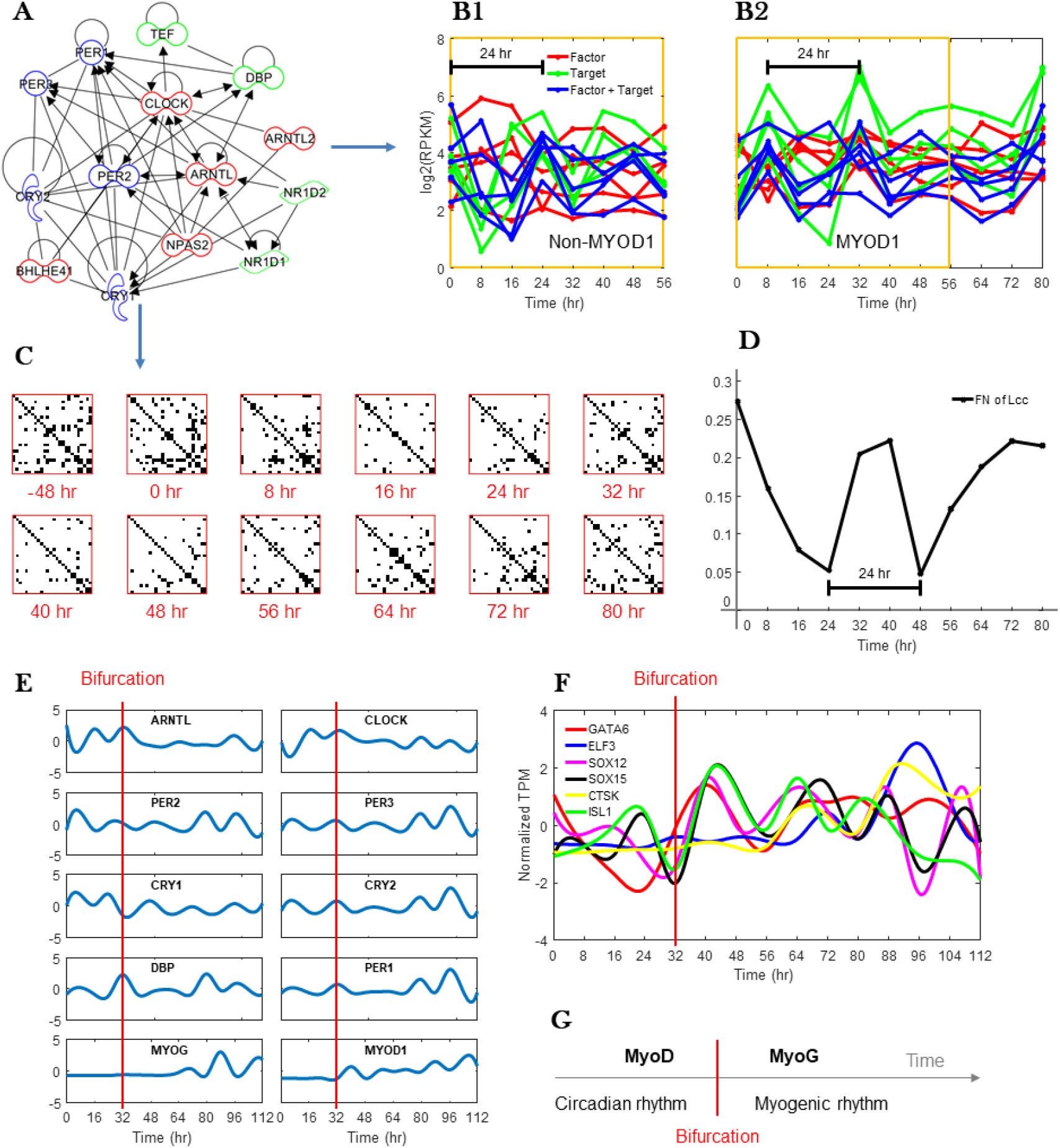
Form and function dynamics of circadian E-box genes. (A) Gene network interactions between circadian E-box genes, derived from Ingenuity Pathway Analysis. (B) Core circadian gene expression over time. (B1) Dexamethasone synchronization. (B2) L-MYOD1 synchronization. Target and factor correspond to genes with E-box targets and TFs that bind to E-box genes, respectively. (C) Hi-C contacts between 26 core circadian genes over time (See Table S2). Rows and columns correspond to core circadian genes, contacts are binary (i.e. any contact between genes at a given time are shown). (D) Network connectivity of the largest connected component (LCC; STAR Methods) of the studied Hi-C contact maps at different time points. (E) Normalized gene expression (FPKM, cubic spline) highlighting oscillation dampening post-bifurcation and post-differentiation medium for select core circadian genes; MYOD1 and MYOG also shown. (F) Normalized transcripts per million (TPM) of transcription factors that are targeted by *MYOG* or *MYOD1 (ELF3*) and only oscillated post-bifurcation point (STAR Methods). (G) Conceptual diagram of biological rhythm entrainment during MYOD1-mediated reprogramming.

Interestingly, the subset of transcripts with oscillatory behavior was different before and after the 32 hr bifurcation point. Endogenous *MYOD1* and *MYOG* expression began close to the bifurcation point and both transcripts displayed oscillatory expression. Additionally, circadian transcript oscillations dampened at 40 hrs, coinciding with the switch to low-serum differentiation medium (Figure 6E). To determine which newly oscillating transcripts were potential targets of MYOD1 and MYOG, we further investigated which transcripts have MYOD1 or MYOG binding motifs in their promoters using MotifMap (Daily et al., 2011), and which were synchronized in expression with MYOD1 and MYOG. Among the oscillating transcripts that fit these criteria, we found six TFs that were oscillatory only after the bifurcation point, have upstream MYOG binding sites, and were synchronized in expression with MYOG. Of these six TFs, only *ELF3* was found to have binding motifs for and synchronized expression with *MYOD1* (Figure 6F). Several of the six oscillatory TFs targeted by MYOG or MYOD1 have been shown to be related to cell differentiation in the literature. Numerous studies have implicated the important role of SOX15 in muscle differentiation (Meeson et al., 2007). GATA6 has been shown to regulate vascular smooth muscle development in several studies (Xie et al., 2015). ISL1 has been shown to interact with CITED2 to induce cardiac cell differentiation in mouse embryonic cells (Pacheco-Leyva et al., 2016). ELF3 is found to play a diverse role in several types of cell differentiation (Böck et al., 2014).

Robust synchronization in expression of circadian genes that are downstream targets of MYOD1 suggests MYOD1 feedback onto circadian gene circuits. After the bifurcation, MYOG was associated with synchronized expression of a subset of important myogenic TFs. These findings support roles for MYOD1 and MYOG in entraining circadian and cell type-specific biological rhythms.

### Changes in Regulatory Elements During Reprogramming

Here we highlight our analyses of early-stage chromatin remodelling gene expression dynamics, super enhancer dynamics, and microRNAs expression.

Examination of early stage RNA-seq data [-48, 16] (hrs) revealed endogenous mechanisms relevant to *MYOD1* transcriptional activation including muscle stage-specific markers and chromatin remodeling factors (See Figure S7). At the 16-hr time point, the combined up-regulation of *DES, MYL·į, TNNT1* and *TNNT2* suggested a differentiation from a myoblast lineage to a skeletal muscle (Gard and Lazarides, 1980; Schiaffino et al., 2015). *EZH2* has been associated with both safe-guarding the transcriptional identity of skeletal muscle stem cells and with terminal differentiation of myoblasts into mature muscle (Juan et al., 2011). *ARID5A*, a regulator of the myotube BAF47 chromatin remodeling complex, is significantly upregulated at 8 hours (P = 7.2 × 10^−5^) and may act to enhance MYOD1 binding to target promoters (Joliot et al., 2014). *NRĄÅ3, MEF2D, SIXĄ, SIX1*, and *SOXĄ* expression are also increased at 8 hr, all of which have important regulatory functions during differentiation in the myogenic lineage (Bentzinger et al., 2012; Ferrán et al., 2016; Jang et al., 2015).

We also investigated how muscle-related super enhancer-promoter (SE-P) interactions change over time throughout MYOD1-mediated reprogramming. To capture these dynamics, we extracted the Hi-C contact between skeletal muscle super enhancer regions and associated genes transcription start site (TSS) (±1kb), as determined by (Hnisz et al., 2013) (618 SE-P regions; STAR Methods). We observed that for these skeletal muscle SE-P Hi-C regions, the strongest amount of contact occured relatively early in the reprogramming process, peaking 16-24 hrs post-L-MYOD1 addition to the nucleus (Figure S8). Exact SE-P contact vs function trends were variable, but a number of important myogenesis genes, such as *TNNI1, MYLPF, ÅCTN2*, and *TNNT3* show strong upregulation in function over time, with an increase in SE-P contact post-MYOD1 activation. Contact vs. function trends for the top 36 upregulated genes are shown in Figure S8 (STAR Methods).

We measured the abundance of 2588 microRNA (miRNA) species with reads from small RNA-seq. Using the edgeR software (Robinson et al., 2010) for data analysis, we identified 266 miRNA species that were significantly up- or down-regulated in expression levels over the time course relative to the baseline control (FDR < 0.05) (Table S6). Among these significant miRNAs, miR-1-3p, miR-133a-3p, and miR-206 have been previously identified as myogenic factor-regulated, muscle-specific species (McCarthy, 2011; Rao et al., 2006). We observed that the three miRNAs, plus miR-133b (FDR = 0.09), significantly increased in expression levels after 4-OHT treatment (Figure S9). Their expression patterns were highly similar to that observed in mouse C2C12 cell differentiation (Rao et al., 2006). The observation of muscle-specific miRNAs, particularly miR-206, which had 1000 fold greater expression at later time points than baseline [mention of muscle specificity redundant here with the above, is McCarthy 2008 needed?] (Figure S9), further supports MYOD1-mediated reprogramming of fibroblasts to myotubes. Notably, the cardiac-specific species miR-1-5p, miR-208a, and miR-208b (McCarthy, 2011) were not detected in our samples.

## DISCUSSION

In this study, we provide analysis on MYOD1-mediated reprogramming of human fibroblasts into the myogenic lineage from a dynamic network perspective. Distinct from previous studies on cellular reprogramming, we generated an enriched time-series dataset integrated with Hi-C, RNA-seq, miRNA, and proteomics data. This provides a comprehensive genome-wide form-function description over time, and allows us to detect early stage cell-fate commitment changes during cellular reprogramming. The discovery of a bifurcation event during cellular reprogramming is exciting because it reveals global and local phenomena associated with the point of transition between cell identities. Our results show that key genes have local dynamics that mirror the global bifurcation. Capturing these dynamics may help us identify genes that are key players in other reprogramming settings, and develop a more universal understanding of the process and requirements for reprogramming between any two cell types.

Our data suggest a direct pathway of reprogramming from fibroblasts to myotubes that bypasses a myoblast intermediate and is associated with expression of *MYOD1* and *MYOG*, but not *MYF5*. Related results have been described in studies on control of the cell cycle during muscle development, in which *MYOD1* and *MYF5* are involved in determination of myogenic cell fate, with a switch from *MYF5* to *MYOG* during muscle cell differentiation (Singh and Dilworth, 2013; Zeng et al., 2016). Moreover, it was theorized in (Del Vecchio et al., 2017) that a reprogrammed bio-system with positive perturbation (i.e. overexpression of one or more specific TFs like MYOD1) would bypass the intermediate state and move directly towards the terminally differentiated state. This claim is consistent with our finding, where the intermediate and terminally differentiated states correspond to myoblast and myotube stages, respectively.

A number of studies have explored the link between *MYOD1* and circadian genes *ARNTL* and *CLOCK*, revealing that ARNTL and CLOCK bind to the core enhancer of the *MYOD1* promoter and subsequently induce rhythmic expression of *MYOD1* (Andrews et al., 2010; Zhang et al., 2012). We found that upon introduction of L-MYOD1, the population of cells exhibits robust synchronization in circadian E-box gene expression. Among these E-box targets are the *PER* and *CRY* gene family, whose protein products are known to repress CLOCK-ARNTL function, thus repressing their own transcription. Additionally, E-box target NR1D1, which is synchronized upon addition of L-MYOD1, competes with ROR proteins to repress *ARNTL* transcription directly. This adds another gene network connection under MYOD1 influence, indirectly acting to repress *ARNTL*, leading us to posit that MYOD1 can affect CLOCK-ARNTL function through E-Box elements, in addition to CLOCK-ARNTL’s established activation effect on MYOD1. Furthermore, these oscillations dampen post-bifurcation point, after which MYOG entrains the oscillations of a distinct subset of myogenic TFs. Therefore, MYOD1-mediated reprogramming and circadian synchronization are mutually coupled, as is the case in many other studies of the reprogramming of cell fate (Umemura et al., 2014; Wagner et al., 2014).

Our proposed bio- and computational technologies shed light on the hypothesis that nuclear reorganization occurs at the time of cell specification and both precedes and facilitates activation of the transcriptional program associated with differentiation (or reprogramming), i.e. form precedes function (Rajapakse and Groudine, 2011). The alternative hypothesis is that function precedes form, that is nuclear reorganization occurs as a consequence of differential transcription and is a consequence of, rather than a regulator of, differentiation programs (Kosak and Groudine, 2004). Our findings support that nuclear reorganization occurs prior to gene transcription during cellular reprogramming, i.e., form precedes function, and that dynamical nuclear reorganization plays a key role in defining cell identity. Our data do not establish a causal relationship, and for this, additional experiments will be necessary. For example, Hi-C and RNA-seq can be supplemented using MYOD1 ChIP-seq to identify the regions of greatest adjacency differences between cell types that correlate with transcription and/or MYOD1 binding.

As demonstrated by our study, network centrality-based analysis allows us to study Hi-C structure from multiple views, and facilitates quantitative integration with gene expression. Accordingly, the detailed connections between network structure and network function in the context of the genome can be used to probe genomic reorganization during normal and abnormal cell differentiation. It will also be helpful to determine whether nuclear architectural remodeling can be both temporally and molecularly separated from transcriptional regulation. Identifying an architectural function for TFs that is distinct from transcription would define a new molecular function with as yet an unknown role in developmental and cancer cell biology. We believe that investigating form-function dynamics will be key in understanding and advancing cell reprogramming strategies. This perspective may have broad translational impact spanning cancer cell biology and regenerative medicine.

## AUTHOR CONTRIBUTIONS

I. R. conceived and supervised the study. H. C., S. R., W. M. and I. R. designed and performed the experiments. S. L., P.-Y. C., A. H. and I. R. contributed network centrality based analytic tools. S. L., H. C., S. R., L. S., G. H and I. R. performed computational analyses and interpreted the data. All authors participated in the discussion of the results. S. L., H. C., S. R., L. A. M. and I. R. prepared the manuscript with input from all authors.

## ACKNOWLEDGMENTS

We thank the University of Michigan Sequencing Core, and especially Jeanne Geskes, for assistance. We thank John Hogenesch for helpful discussions on circadian rhythms. We thank Daniel Burns and Stephen Lindsly for critical reading of the manuscript and helpful discussions. We extend special thanks to James Gimlett and Srikanta Kumar at Defense Advanced Research Projects Agency (DARPA) for support and encouragement. This work is supported, in part, by the DARPA Biochronicity Program and the DARPA Deep-Purple and FunCC Program. We also acknowledge the seminal work of Mark Groudine and late Hal Weintraub, whose ideas continue to guide our thinking.

## SUPPLEMENTAL FIGURE TITLES AND LEGENDS

**Figure S1:**
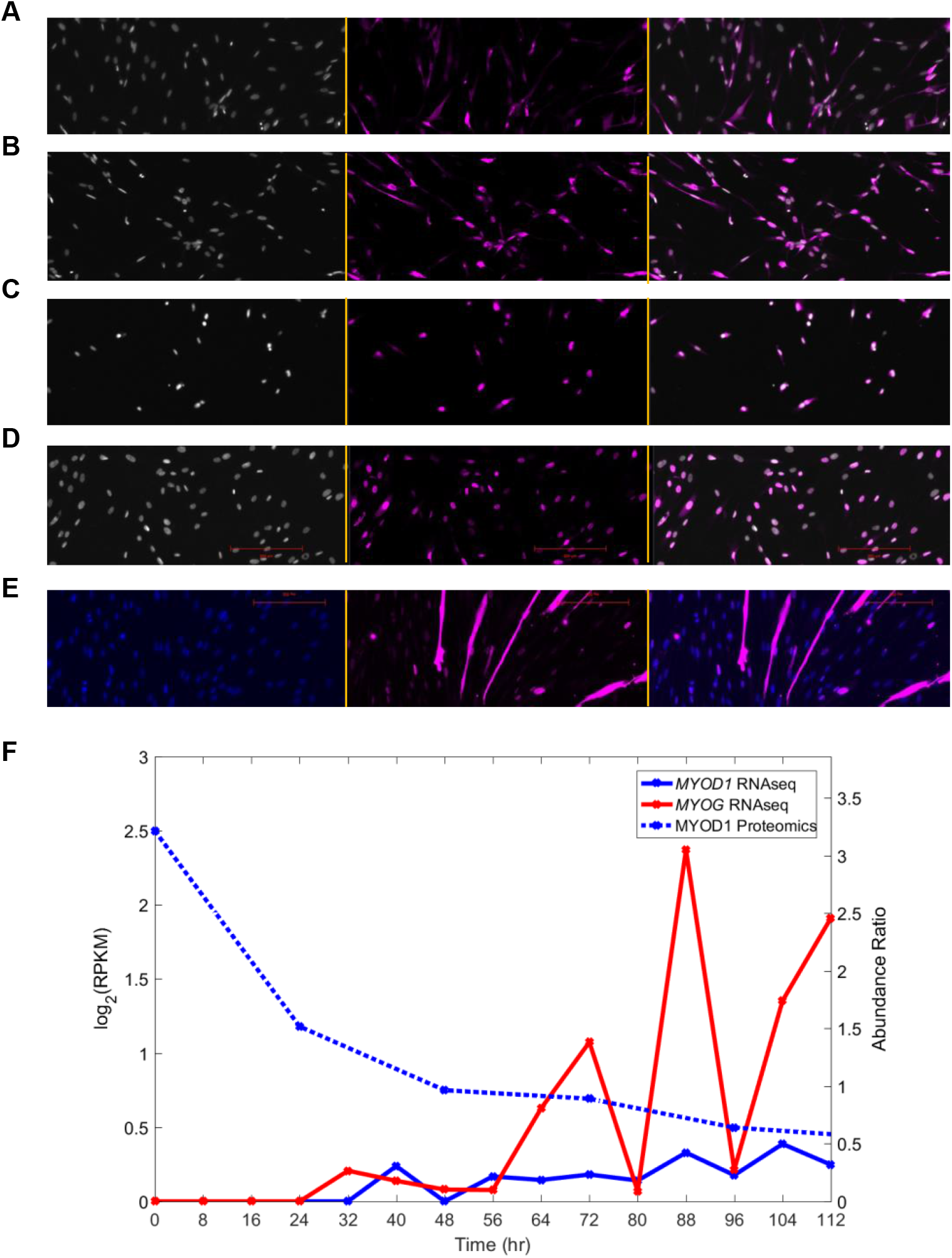
Fluorescent micrographs showing immunocytochemistry dynamics of MYOD1 localization and the detection of MYH1 expression, and quantification of RNA and protein abundance; Related to Figure 1. (A-E) Left panels show DAPI staining of nuclei, middle panels show anti-MYOD1 staining (A-D) and anti-MYH1 staining (E), right panels are overlay of left and middle images, respectively. (F) Time-series RNA-seq (solid line) and Proteomic (dashed line) quantification of RNA and protein abundance, respectively, for MYOD1 (blue) and MYOG (red).

**Figure S2:**
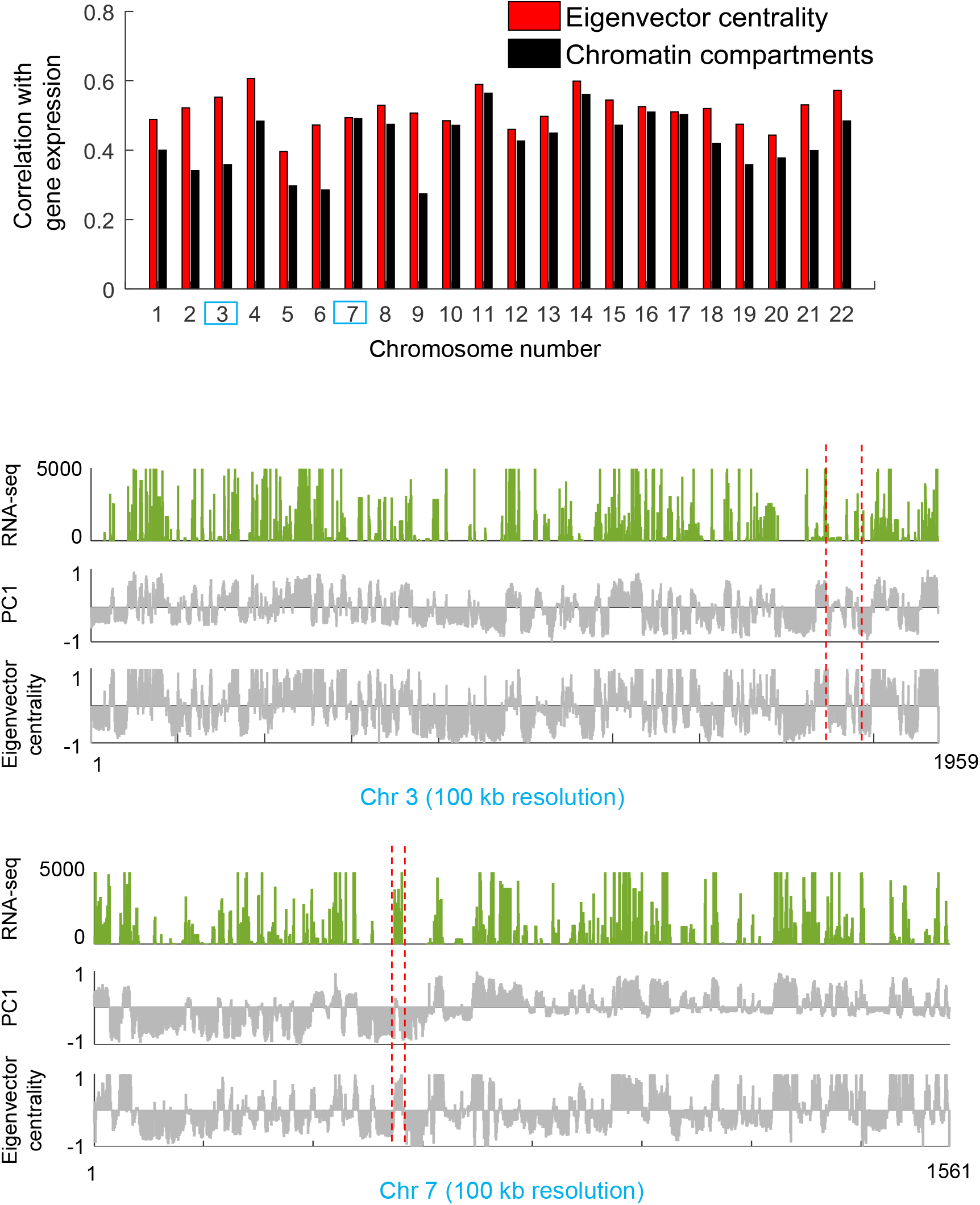
Eigenvector centrality refines active and inactive chromatin domains; Related to Figure 2 and 3. Eigenvector centrality yields a higher correlation with gene expression than chromatin partition conventionally defined by the first principal component (PC1) of the spatial correlation matrix of Hi-C data (Lieberman-Aiden et al., 2009). Chromosome 3 and 7 are shown as examples.

**Figure S3:**
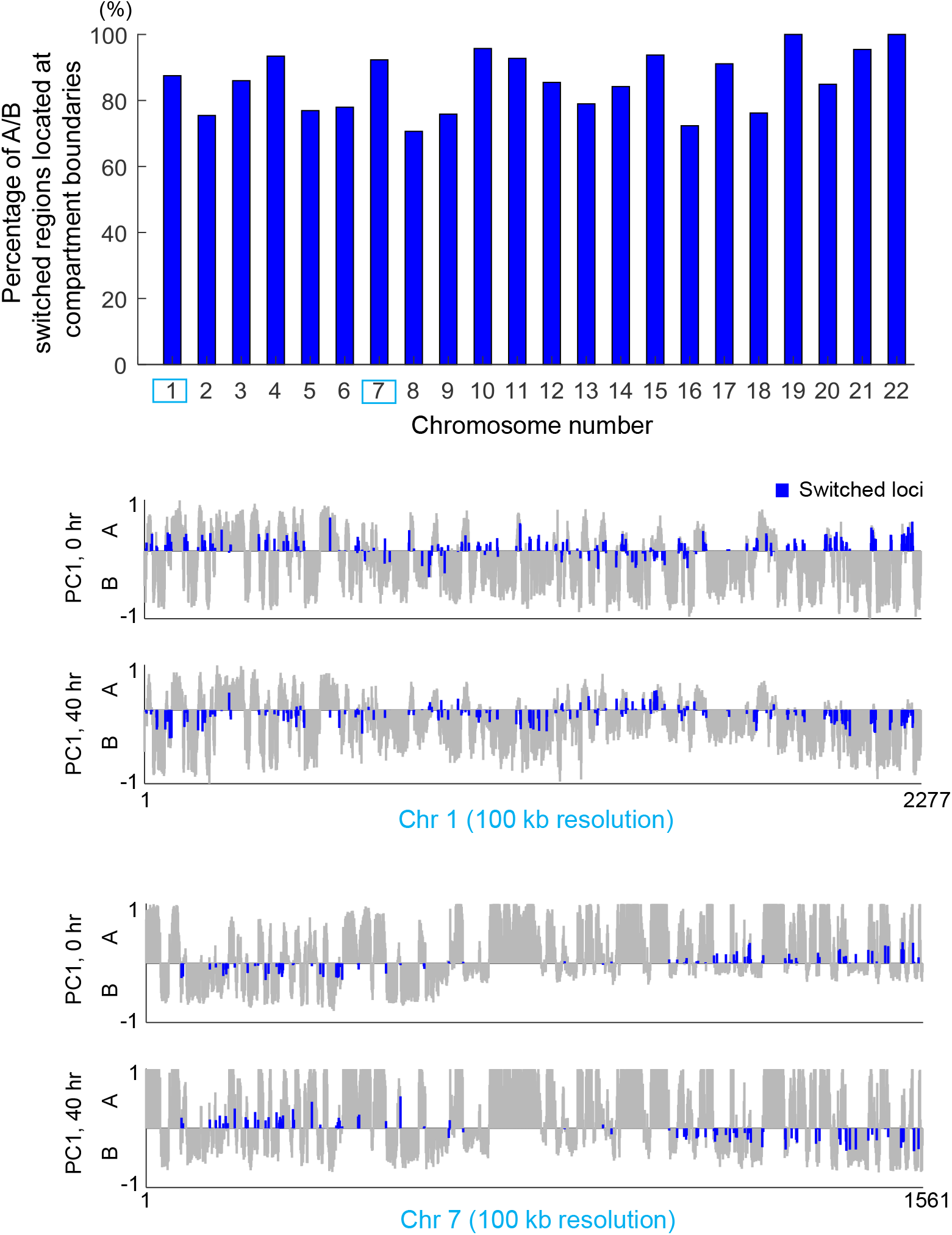
Chromatin compartment change appears at their boundary regions; Related to Figure 3. Over 70% A/B switched bins are at A/B boundary loci. Chromosome 1 and 7 are shown as examples of chromatin compartment switching from 0 to 40 hrs.

**Figure S4:**
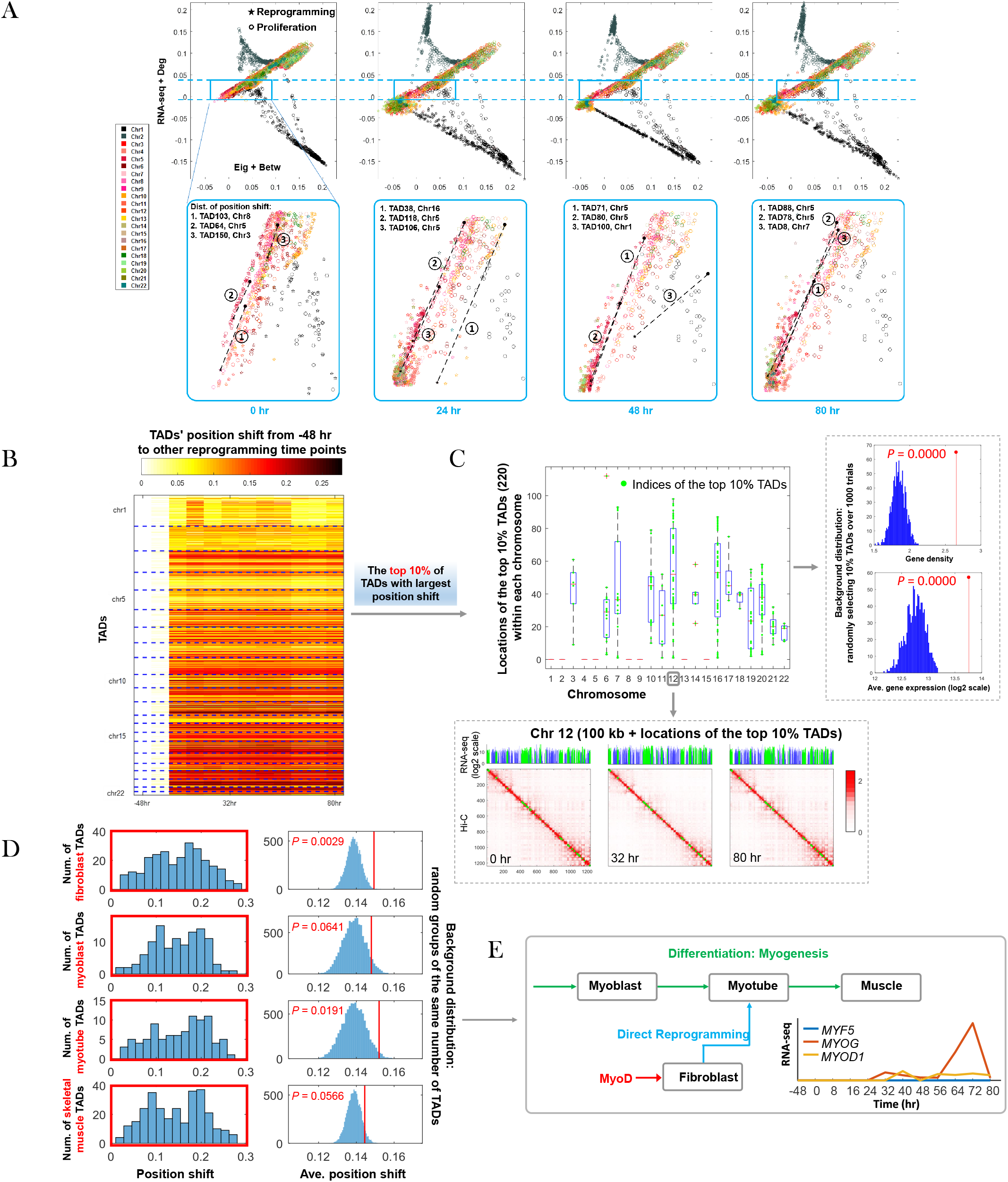
A direct pathway from fibroblasts to myotubes; Related to Figure 2. (A) 2D representations of TAD-scale form-function features at time 0, 24, 48 and 80 hrs. The star marker represents the coordinate of a TAD at the reprogramming time instant. The circle marker represents the TAD at the stage of fibroblast proliferation (-48 hr). A specified region of data configuration (top plots) is magnified in bottom plots, where three topologically associating domains (TADs) with the 1st, 10th and 20th largest position shift (from proliferation to reprogramming) are marked. (B) Heatmap of TADs’ position shift from –48 hr to reprogramming time points. (C) TADs with top 10% largest position shift. *Top left*: Locations of the identified TADs over chromosomes. *Bottom left*: Example of identified TADs (green color) at chromosome 12 (100kb-binned Hi-C) together with gene expression at time 0, 32 and 80 hrs. *Right*: P values of gene density and average gene expression. (D) Position shift of TADs that involve fibroblast, myoblast, myotube, and skeletal muscle related genes, respectively. *Left*: Histograms of TADs’ position shift for each gene module of interest. *Right*: P value of average position shift for each gene module.(E) Direct pathway from fibroblasts to myotubes evidenced by gene expression of three myogenic regulatory factors: *MYF5, MYOD1* and *MYOG*.

**Figure S5:**
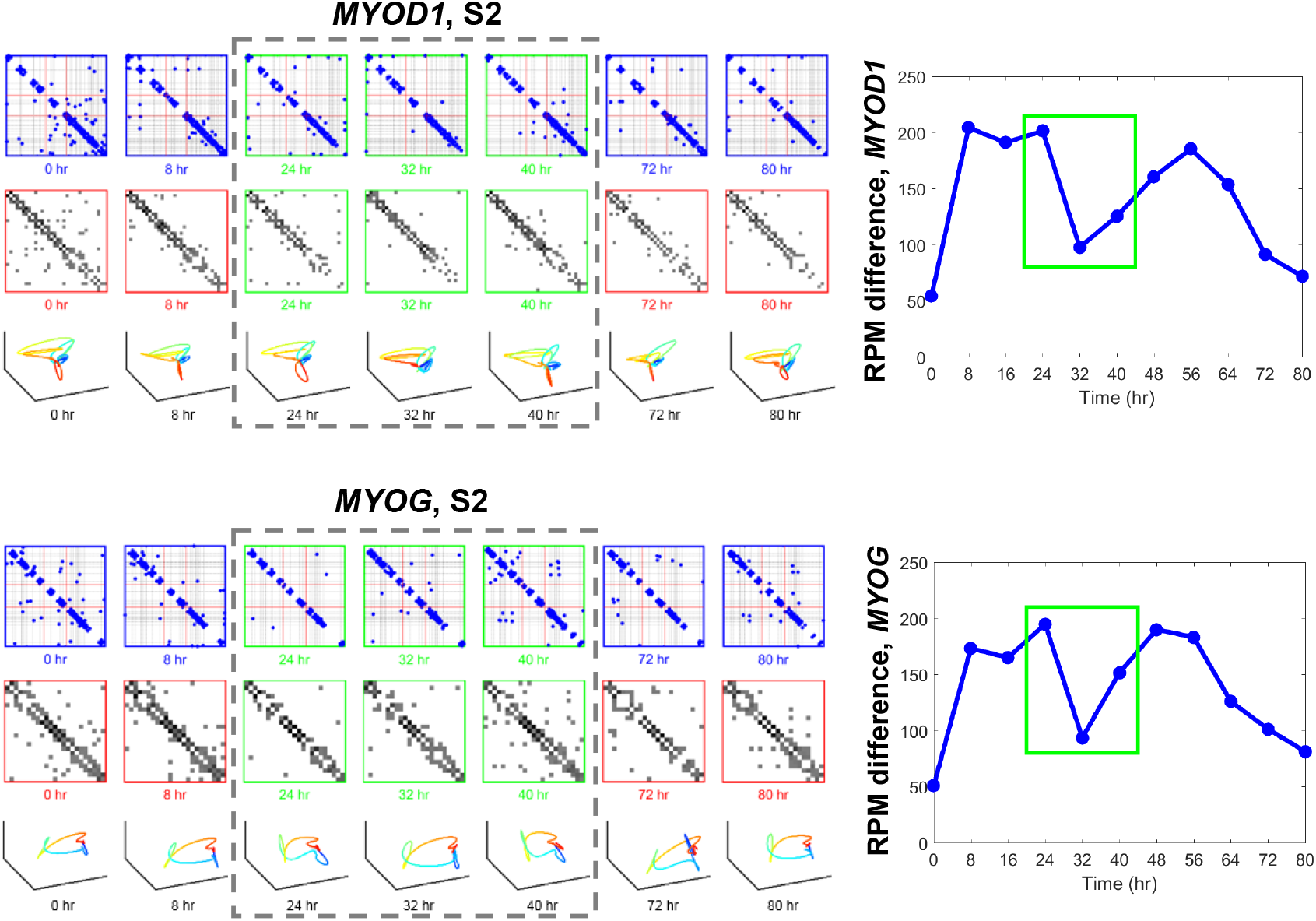
Genomic dynamics of *MYOD1* and *MYOG*; Related to Figure 2. *Top left* or *Bottom left*: First row depicts Hi-C contact maps of *MYOD1* (or *MYOG*) at base pair scale, where blue points are contacts, red lines depict gene boundaries, and dashed black lines depict MboI cut-sites. Middle rows show Hi-C matrices binned by MboI cut sites and normalized by RPM. Bottom row shows 3D gene models, given by cubic Bézier curves that fits 3D representation of MboI binned contact matrices using Laplacian eigenmaps (STAR Methods). *Top right* or *Bottom right*: Summation of entry-wise differences of Hi-C matrices for *MYOD1* (or *MYOG*) between time points.

**Figure S6:**
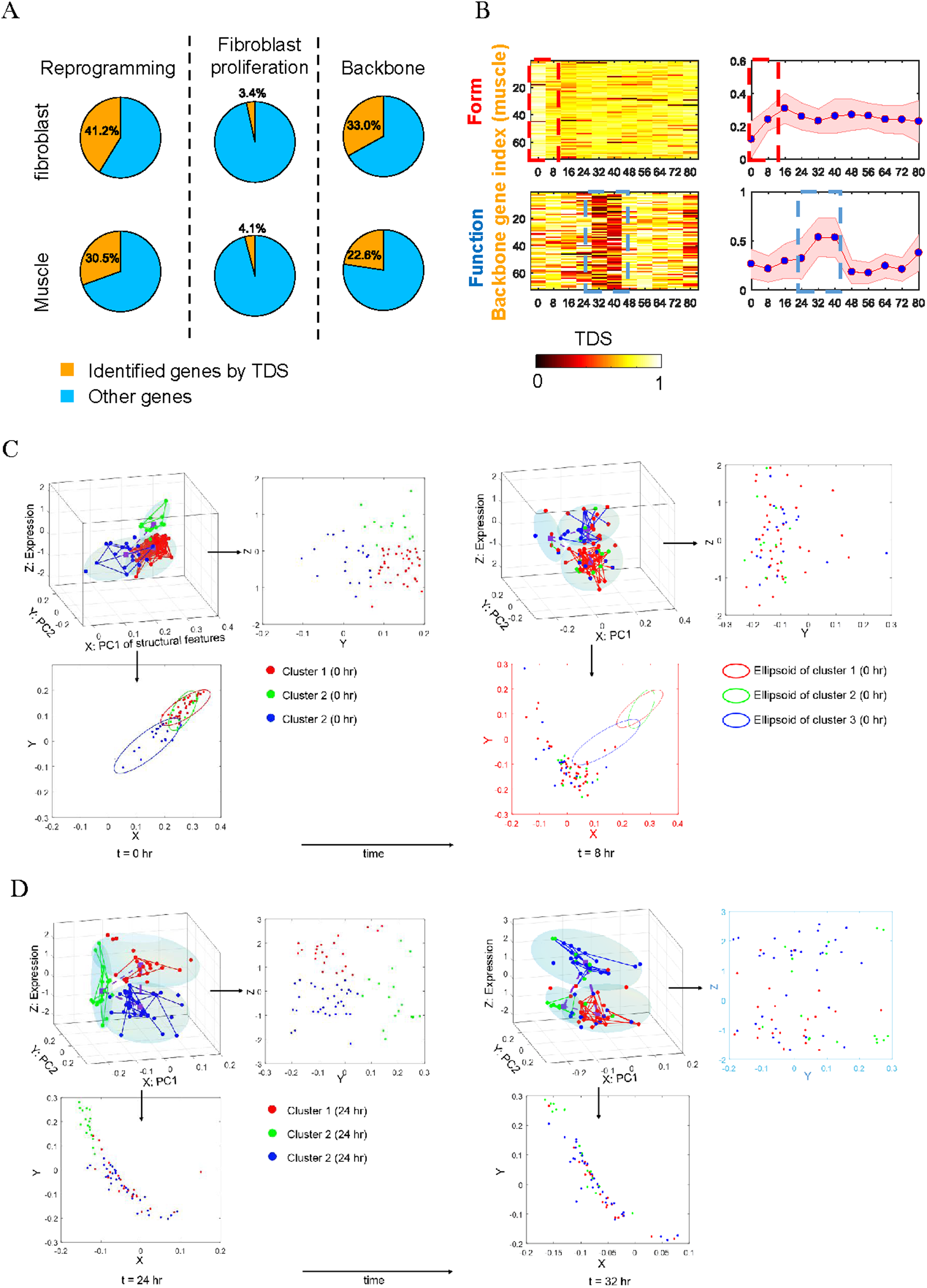
Backbone genes in fibroblast and muscle gene module; Related to Figure 4. (A) Pie charts revealing the portion of backbone genes within each gene module. *Left*: Portion of genes recognized by form-function TDS during cellular reprogramming. *Middle*: Portion of the aforementioned genes that are also active during fibroblast proliferation. *Right*: Backbone genes given by the set of genes extracted from reprogramming but excluding those from proliferation. (B) Heatmap of form and function TDS for muscle-related backbone genes. (C) 3D configuration of muscle-related backbone genes in form-function space from 0 to 8 hrs, highlighting significant form change. The edge represents Hi-C contact between genes. Three clusters of genes at 0 hr are marked by red, green, and blue, respectively. The 3D ellipsoid determined by MVE provides the clustering envelope at the current time, where its centroid is marked by purple square. (D) 3D configuration of muscle-related backbone genes in form-function space from 24 to 32 hrs, highlighting significant function change.

**Figure S7:**
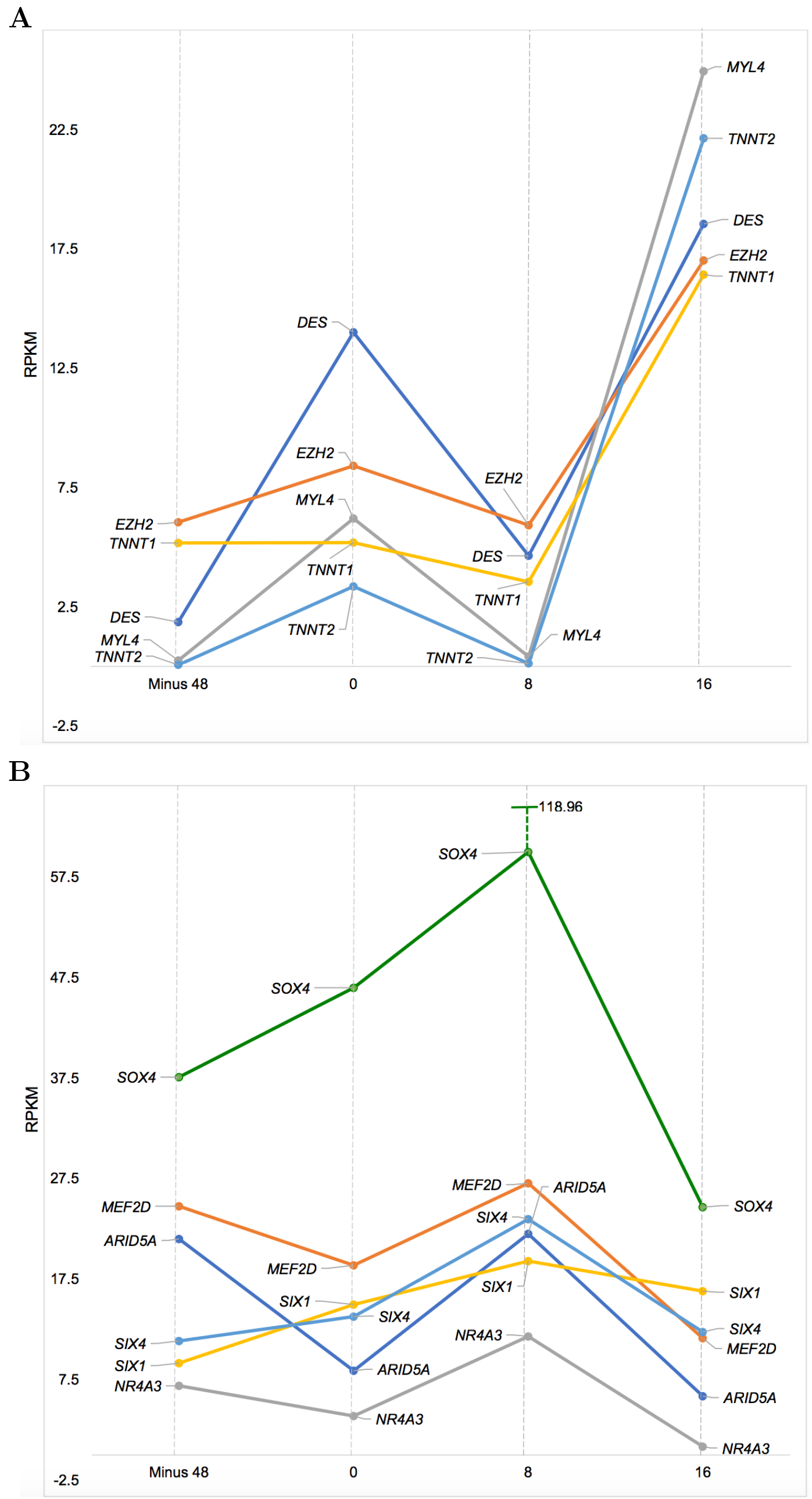
Early-phase expression dynamics of genes related to muscle cell terminal differentiation and chromatin remodeling. (A) Genes encoding proteins involved in adult muscle function, including components of the contractile apparatus (DES, MYL4, TNNT1, TNN2), and EZH2, which is a repressor and is involved in myogenesis. (B) Chromatin remodeling factors and master transcription factors act cooperatively with MYOD1 to drive proliferating human fibroblasts into muscle cells. They include ARID5A, part of the BAF47 muscle remodeling complex which acts in cooperation with MYOD1, MEF2D, which drives differentiation of myotubes to skeletal and cardiac muscle, NR4A3 (aka NOR1) involved in differentiation of myotubes into smooth muscle, and SIX1, SIX4 and SOX4, which control the differentiation of myotubes into muscle cells.

**Figure S8:**
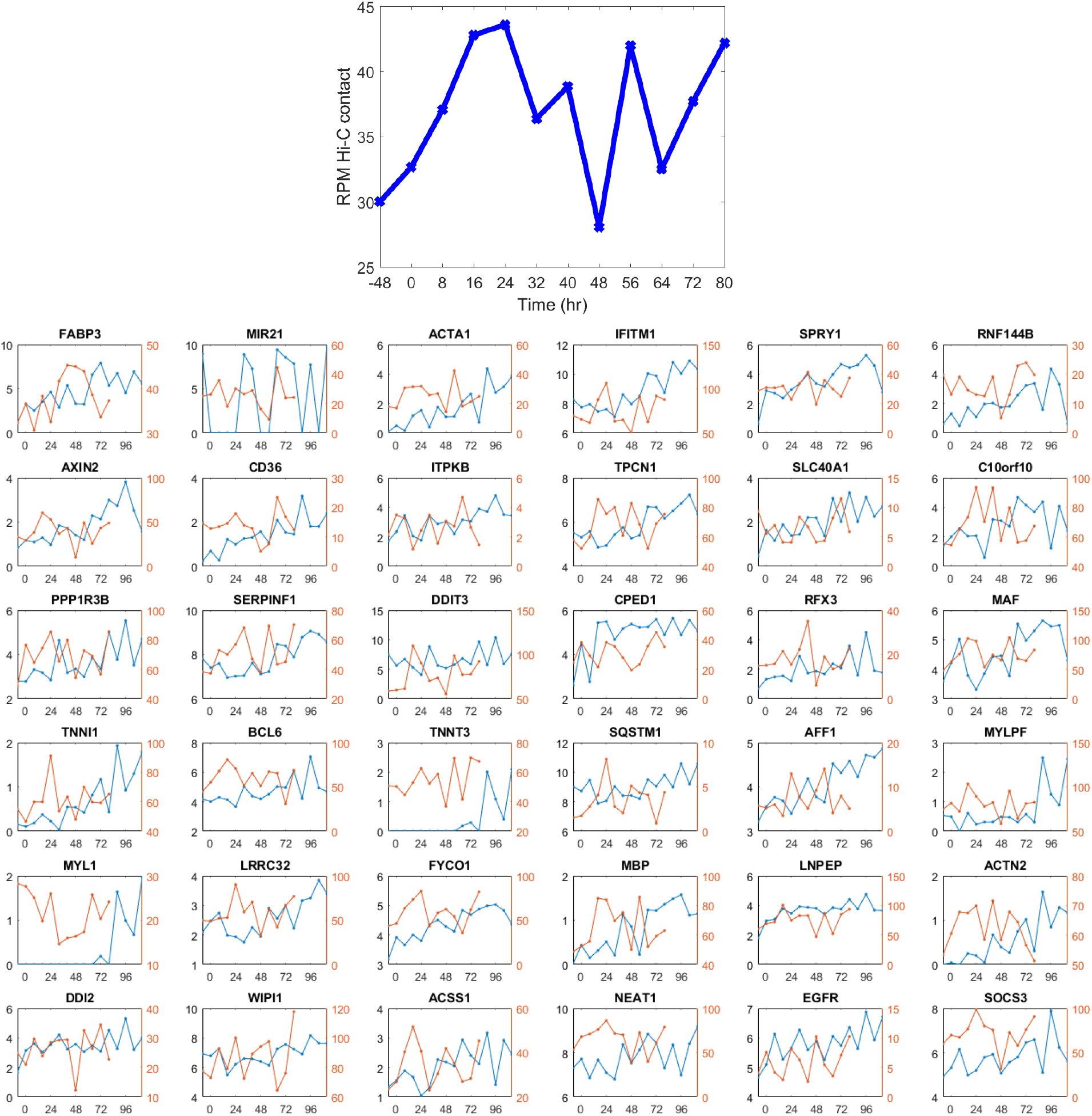
Form and function of super enhancers and associated genes over time. *Top*: Average Hi-C RPM contact between Potential super enhancer and associated gene TSS regions over time, as defined by Hnisz et al. (2013). *Bottom*: Top upregulated SE-P genes, log2(FPKM) (Blue) and SE-P Hi-C normalized contact (Red; see STAR Methods) over time.

**Figure S9:**
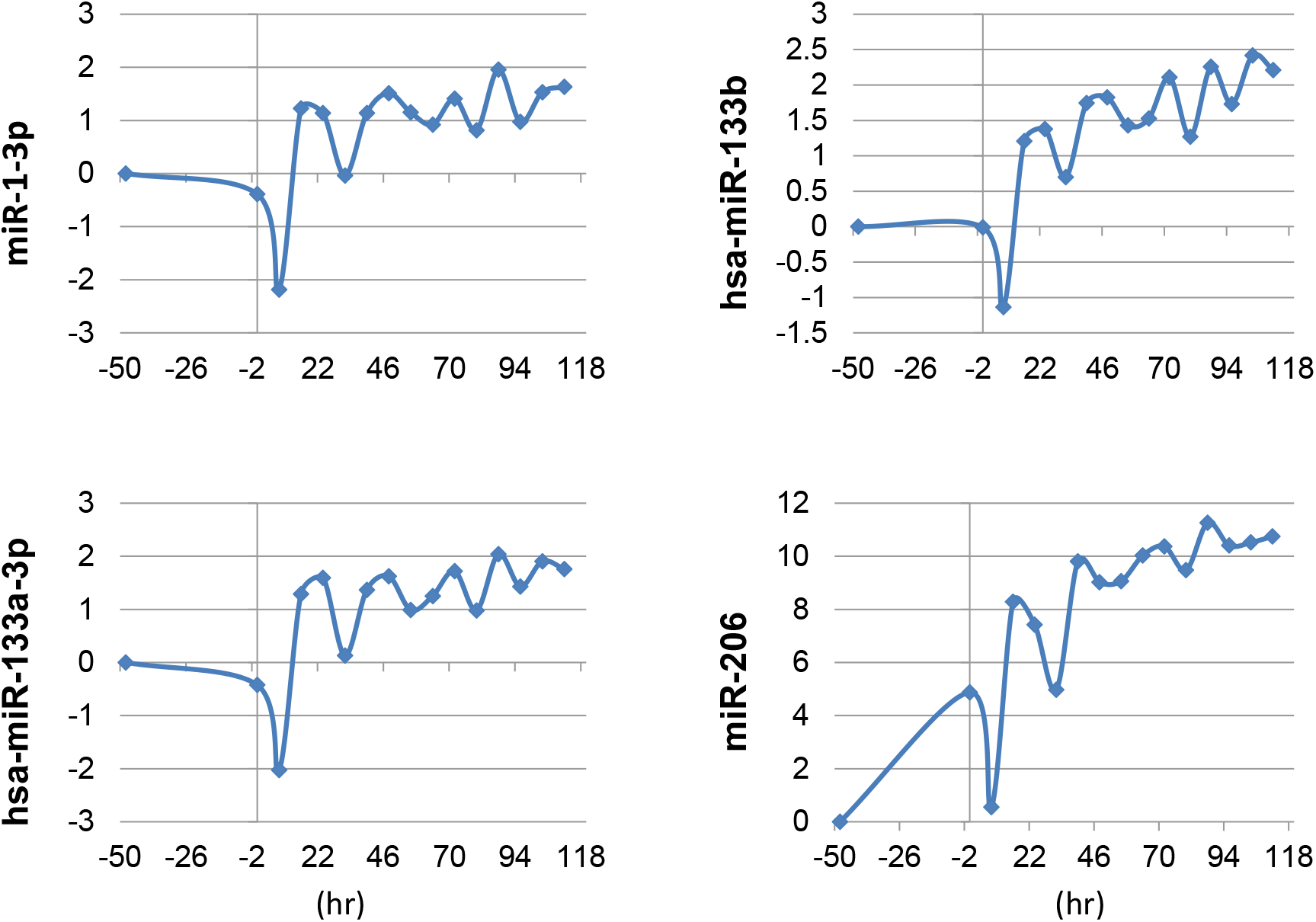
Four muscle specific miRNAs significantly increased expression levels in the later time points relative to the baseline control. X-axis corresponds to sampling time points, and y-axis shows log-scale differences at other time points compared to baseline (-48 hrs).

## STAR**⋆** METHODS

The outline of detailed methods:

- KEY RESOURCES TABLE
- CONTACT FOR REAGENT AND RESOURCE SHARING
- METHOD DETAILS

∘ Generation of a human *MYOD1* expressing construct
∘ Cell culture, lenti-viral transduction, and induction of *MYOD1*
∘ RNA-seq and small RNA-seq
∘ Crosslinking of cells for Hi-C
∘ Generation of Hi-C libraries for sequencing
∘ Generation of Hi-C matrices
∘ Reverse transcriptional polymerase chain reaction (RT-PCR) analysis
∘ Immunocytochemistry analysis
- QUANTIFICATION AND STATISTICAL ANALYSIS

∘ Scale-adaptive gene expression
∘ Scale-adaptive Hi-C matrix
∘ Network representation of 4DN: graph Laplacian and Fiedler number
∘ Structural feature extraction via network centrality measures
∘ Integration of form and function
∘ Data representation on low-dimensional non-linear manifolds
∘ Fitting the data: minimum volume ellipsoid
∘ Temporal difference score (TDS)
∘ A/B compartment switching analysis
∘ Bifurcation: branching trajectory and statistical significance
∘ Bifurcation identification at gene level
∘ Identification of genes of interest
∘ Statistical significance of TDS of genes
∘ Identification of MYOD/MYOG mediated oscillatory gene expression
∘ Super enhancer-promoter region dynamics
- DATA AND SOFTWARE AVAILABILITY

## KEY RESOURCES TABLE

Please see a submitted separate table named by ‘Key resource table’.

## CONTACT FOR REAGENT AND RESOURCE SHARING

Further information and requests for resources and reagents should be directed to and will be fulfilled by the Corresponding Contact, Indika Rajapakse.

## METHOD DETAILS

### Generation of a human MYOD1 expressing construct

We generated a lenti-construct (lenti-hMYOD1-mER(T)) expressing the human myogenic differentiation factor 1 protein (hMYOD1) fused with a tamoxifen-specific binding domain (mER(T)) derived from mouse estrogen receptor 1 (Kimura et al., 2008). The open reading frame (ORF) for the fusion protein was synthesized at IDT (Integrated DNA technologies) as one gBLOCK, and cloned into the NheI/EcoRI sites of a lenti-vector (obtained from the University of Michigan Vector Core). The expression of the fusion protein is driven by a CMV promoter. The lenti-viral particles were produced at the University of Michigan Vector Core facility for transduction of human BJ fibroblasts with normal karyotype (Cat# CRL2522, ATCC).

### Cell culture, lenti-viral transduction, and induction of MYOD1 reprogramming

BJ cells were propagated in growth medium (GM) composed of DMEM (Cat# 11960069, Thermo Fisher Scientific), 10% fetal bovine serum (Cat# 10437028, Thermo Fisher Scientific), 1x non-essential amino acids (Cat#11140050, Thermo Fisher Scientific), and 1x Glutamax (Cat# 35050061, Thermo Fisher Scientific). The day before viral transductions, fibroblasts at the 7th passage were plated in 6-well plates or T75 flasks in 13 mL of GM. We plated 1 × 10^5^ cells per well in 6-well plates for RNA extraction, and 2 × 10^6^ cells per flask T75 flasks for Hi-C and proteomics sampling. The cells were incubated in an incubator at 37° C with 5% of CO2.

Lenti-viral transduction was performed the next day after plating the cells. We used a MOI (multiplicity of infection) of 15 to transduce the cells in 8 mL GM plus 4 μg/mL of polybrene (Cat# 107689, Sigma-Aldrich). The transduction incubation was carried out in an incubator at 37° C with 5% CO2 for 12 hours. After the incubation, the transduction medium was removed, and the cells were washed with PBS (Cat# 10010049, Thermo Fisher Scientific), then fed with 13 mL of fresh GM to continue incubation for 24 hours.

To induce MYOD1 into the nucleus for myogenic reprogramming, we treated the cells transduced with lenti-hMYOD1-mER(T) by adding (Z)-4-Hydroxytamoxifen (4-OHT) (Cat# H7904, Sigma-Aldrich) to a final concentration of 1 μM to each flask in GM for two days. This treatment translocated the hMYOD1-mER(T) protein in the cytoplasm to the nucleus for MYOD1-meditated myogenic reprogramming (Kimura et al., 2008). To induce differentiation after 4-OHT treatment, we washed the cells twice with PBS, and changed to differentiation medium consisting of DMEM supplemented with 2% horse serum (Kimura et al., 2008).

### Crosslinking of cells for Hi-C

During a time course sampling, at each time point the cells in a T75 flask were washed with 10 mL PBS, and then incubated with 15 mL of 1% formaldehyde prepared in PBS at room temperature for exactly 10 min. To quench the crosslinking reaction, glycine of 2.5 M was added to the flask to a final concentration of 0.2 M, and incubated for 5 min at room temperature on a rocking platform, then on ice for at least 15 min to stop crosslinking completely. The cells were scraped off with a scraper and transferred into 15 mL tubes. The crosslinked cells were collect with centrifugation at 800 x g for 10 min at 4° C. The cells collected were washed in 1 mL ice-cold PBS briefly, and spun down at 800 x g for 10 min at 4^°^ C. After centrifugation, the supernatant was discarded completely, and the cells were snap-frozen in liquid nitrogen and stored at −80° C for Hi-C library construction.

### RNA-seq and small RNA-seq

We used the miRNeasy Mini Kit (Cat# 217004, Qiagen) for total RNA isolation according to the manufacturers manual. The RNA samples extracted from each sampling time point were treated with RNase-Free DNAase I (Cat# 79254, Qiagen) to clean up any DNA contamination.

All RNA-seq and small RNA-seq data were generated at the University of Michigan Sequencing Core facility. RNA quality control (QC) was performed at the Core. The QC results from the TapeStation analysis (Agilent, Technologies) showed that the samples RNA integrity number (RIN) was > 9.8. The RNA-seq libraries were prepared according to the TruSeq RNA Library Prep Kit v2 chemistry (Cat# RS-122-2001, Illumina). The small RNA-seq libraries were prepared with the NEBNext^®^ Small RNA Library Prep Set for Illumina (Cat# E7330S, New England Biolabs, NEB).

We sequenced the mRNA species for each samples to produce the RNA-seq dataset, and the small RNA species to obtain the miRNA-seq dataset. Sequence reads were generated on the Illumina HiSeq 2500 platform with the V4 single end 50-base cycle. We used an in house pipeline for sequence read QC (FastQC), genome mapping and alignment (Tophat & Bowtie2), and expression quantification (Cufflinks). We used edgeR (Robinson et al., 2010) for differential expression analysis.

### Generation of Hi-C libraries for sequencing

We adapted the in situ Hi-C protocols from Rao et al (Rao et al., 2014) with slight modifications. Briefly, we used 1% formaldehyde for chromatin cross-linking. We used approximately 2.5 × 10^6^ cells for each Hi-C library construction. The chromatin was digested with restriction enzyme (RE) MboI (Cat# R0147M, NEB) overnight at 37° C with rotation. RE fragment ends were filled in and marked with biotin-14-dATP (Cat# 19524016, Thermo Fisher Scientific), and ligated with T4 DNA ligase (NEB, M0202). After the chromatin decross-linking and DNA isolation, DNA samples were sheared on a Covaris S2 sonicator to produce fragments ranging in size of 200-400 bp. The biotinylated DNA fragments were directly pulled down with the MyOne Streptavidin C1 T1 beads (Cat# 65001, Thermo Fisher Scientific). The ends of pulled down DNA fragments repaired, and ligated to indexed Illumina adaptors. The DNA fragments were dissociated from the bead by heating at 98° C for 10 minutes, separated on the magnet, and transferred to a clean tube.

Final amplification of the library was carried out in multiple polymerase chain reactions (PCR) using Illumina PCR primers. The reactions were performed in 25 μL scale consisting of 25 ng of DNA, 2 μL of 2.5mM dNTPs, 0.35 μL of 10 μM each primer, 2.5 μL of 10X PfuUltra buffer, PfuUltra II Fusion DNA polymerase (Cat# 600670, Agilent). The PCR cycle conditions were set to 98° C for 30 seconds as the denaturing step, followed by 14 cycles of 98° C 10 seconds, 65° C for 30 seconds, 72° C for 30 seconds, then with an extension step at 72° C for 7 minutes.

After PCR amplification, the products from the same library were pooled and fragments ranging in size of 300-500 bp were selected with AMPure XP beads. The size selected libraries were sequenced to produce paired-end Hi-C reads on the Illumina HiSeq 2500 platform with the V4 of 125 cycles.

### Generation of Hi-C matrices

We standardized an in house pipeline to process Hi-C sequence data. With this pipeline, FastQC (http://www.bioinformatics.bbsrc.ac.uk/projects/fastqc/) was used for quality control of the raw sequence reads. Paired-end reads with excellent quality were mapped to the reference human genome (HG19) using Bowtie2 (Langmead and Salzberg, 2012), with default parameter settings and the “–very-sensitive-local” preset option, which produced a SAM formatted file for each member of the read pair (R1 and R2). HOMER (http://homer.salk.edu/homer/interactions/) was used to develop the contact matrix with “makeTagDirectory” with the tbp 1 setting, and with “analyzeHiC” with the “-raw” and “-res 1000000” settings to produce the raw contact matrix at 1Mb resolution, or with the “-raw” and “-res 100000 settings to produce contact matrix at 100kb resolution.

### Reverse transcriptional polymerase chain reaction (RT-PCR) analysis

The cDNA templates for RT-PCR were synthesized from 1 μg RNA using the SuperScript^®^ III First-Strand Synthesis System (Cat# 18080051, Thermo Fisher Scientific). Targets amplicons of corresponding genes (see Key Resources Table) were amplified in 20 μL reactions using the following settings: initial denaturation was performed at 95° C for 5 min, followed by 30 cycles at 95° C for 15 seconds, 56° C for 30 seconds, and 72° C for 20 seconds. The PCR reactions were then incubated for a final extension step at 72° C for 5 min. The products were analyzed on 1.5% agarose gel. The gel image was taken on an imaging station (Universal Hood II, Bio Rad).

### Immunocytochemistry analysis

Cells were grown in appropriate media on washed and autoclaved 12mm round 1.5 glass coverslips placed in 12 well culture plates. At harvest, coverslips were rinsed briefly in phosphate-buffered saline pH 7.4 (PBS), treated with 4% paraformaldehyde in PBS for 10 min at room temperature, then washed three times in PBS at 5 minutes per wash. Cells were dehydrated in a series of ice-cold ethanol concentration steps, 50%, 70%, 90% and 100% at 5 minutes per step, and stored at 4° C until staining. Rehydration reversed the concentration series, with two washes in cold PBS at the end. Cells were permeabilized for 10 min in a PBS 0.25% Triton X-100 solution at RT, and then washed in PBS three times for 5 min per wash. Blocking of non-specific antibody binding was performed with 1% BSA PBST (PBS + 0.1% Tween 20) for 30 minutes, followed by immunostaining using primary antibody (DSHB anti-MHC MF20 diluted 1:20, and/or Thermofisher anti-MyoD diluted 1:250) in 1% BSA in PBST in a humidified chamber for 1 hr at room temperature (RT). The primary solution was removed, cells were washed three times in PBS at 5 min per wash, and the fluorescent secondary, Alexa Fluor 594 goat anti-mouse IgG in 1% BSA PBST was applied for 1 hr at RT in the dark. The secondary antibody solution was then removed and the cells were washed three times with PBS for 5 min each in the dark. Cells were mounted on slides with Prolong Gold anti-fade reagent with DAPI, and imaged.

## QUANTIFICATION AND STATISTICAL ANALYSIS

### Scale-adaptive gene expression

Hi-C matrices are commonly created by converting number of interaction reads into values at fixed resolution bins (e.g., 100kb, 1Mb). However, RNA-seq data (FPKM) are generated at gene level. For consistent analysis of form and function, we transform the RNA-seq data at gene level to its counterpart at bin level, namely,

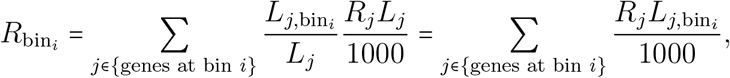

where *L_j_* is the length of gene *j* at base-pair unit, 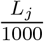 is at kilobase-pair unit, *L*_*j*,bin_. is the length of the portion of gene *j* belonging to bin *i*, *R_j_* signifies the FPKM value at gene *j*, and *R*_bin_*i*__ denotes the total RNA-seq RPM value at bin *i*.

### Scale-adaptive Hi-C matrix

It is expected that nearby loci in linear base-pair distance are more likely to be ligated than distant pairs. This makes a Hi-C matrix highly diagonally dominant and conceals the contact pattern embedded in the matrix. In order to alleviate this effect, we normalize the counts by their contact probability as a function of the linear distance, namely, each entry of the matrix is normalized by its expected contact value (expected-observed method). This is equivalent to normalization of the Hi-C matrix by a Toeplitz structure whose diagonal constants are the mean values calculated along diagonals of the observed matrix; see details in (Chen et al., 2015, SI).

Similar to scale-adaptive gene expression, we are also able to construct gene-resolution contact maps by calculating the contact frequency of two genes, which is normalized by lengths of genes (Chen et al., 2015). Moreover, to construct TAD-scale contact matrices, we begin by normalizing both intra- and inter-chromosome Hi-C matrices at 100kb resolution, and then compute the density of genome contacts among TADs. TAD boundaries here are defined based on (Dixon et al., 2012). Given TADs *i* and *j*, the resulting contact map **T** is given by

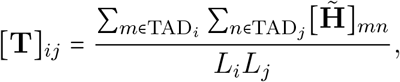

where 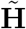 is the normalized Hi-C matrix (100kb-binned Hi-C in our analysis), and *L_i_* is the size of TAD*_i_*. Since the TAD-scale contact matrix is dense, we apply thresholding to convert it to a sparse version by retaining only interactions that exceed the 50th-percentile of Hi-C contact at TAD scale.

### Network representation of 4DN: graph Laplacian and Fiedler number

Let *G_t_* = (*V*, *E_t_*) denote a weighted undirected graph at time *t*, where *V* is a node set with cardinality |*V*| = *n*, and *E_t_* ⊂ {1, 2, …, *n*} × {1, 2, …, *n*} is an edge set at time *t*. The Hi-C matrix **H***_t_* can then be interpreted as an adjacency matrix corresponding to *G_t_*, where (*i*, *j*) ∈ *E_t_* if there exists interactions between node *i* and *j* with edge weight [**H***_t_*]_*ij*_ > 0 and [**H***_t_*]*_ij_* = 0 otherwise. Here nodes represent fixed-size bins, genes or TADs. It is often the case that a graph/network is represented through the graph Laplacian matrix, **L***_t_* = **D***_t_* − **H***_t_*, where **D***_t_* = diag(**H***_t_***1**) is the degree matrix of *G_t_*, **1** denotes the vector of all ones, and diag(**x**) signifies the diagonal matrix with diagonal vector **x**. Given **L***_t_*, the Fiedler number and the Fiedler vector is defined by the second smallest eigenvalue and its corresponding eigenvector. It is known from spectral graph theory (Chung, 1997) that *G_t_* is connected (namely, there exists a path between every pair of distinct nodes) if and only if the Fiedler number is nonzero, and the entrywise signs of Fiedler vector encodes information on network partition. For a network with Fiedler number of being zero, we can extract its largest connected component (LCC), namely, the largest subgraph with nonzero Fiedler number.

### Structural feature extraction via network centrality measures

A network/graph centrality measure is a quantity that evaluates the influence of each node to the network, and thus provides essential topological characteristics of nodes (Newman, 2010). In what follows, we introduce the key centrality measures used in our analysis and elaborate on the rationale behind them.

- Degree. A nodal degree is defined as the sum of edge weights (namely, Hi-C contacts) associated with each node,

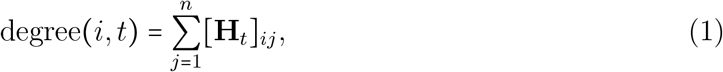

where degree(*i*, *t*) denotes the degree of node *i* at time *t*. We remark that degree(*i*, *t*) exhibits the spatial proximity between node *i* to other nodes.
- Eigenvector centrality. The eigenvector centrality is defined as the principal eigenvector of the adjacency matrix corresponding to its largest eigenvalue, namely

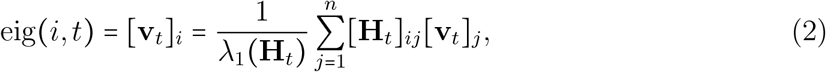

where λ_1_(**H***_t_*) is the maximum eigenvalue of **H***_t_* in magnitude, and **v***_t_* is the associated eigenvector, namely λ_1_(**H***_t_*)**v***_t_* = **H***_t_***v***_t_*. It is clear from (2) that the eigenvector centrality relies on the principle that a node has more influence if it is connected to many nodes which in turn are also considered to be influential. Different from degree centrality, the eigenvector centrality takes the full network topology into account.
- Betweenness. Betweenness is the fraction of the number of shortest paths passing through a node relative to the total number of shortest paths in the connected network. The betweenness of node *i* at time *t* is defined as

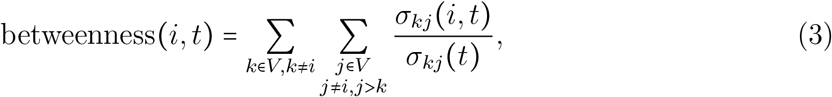

where *σ_kj_*(*t*) is the total number of shortest paths from node *k* to *j* at time *t*, and *σ_kj_*(*i*, *t*) is the number of such shortest paths passing through node *i*. Betweenness characterizes potential hub nodes in the network, and thus a node with high betweenness has the potential to disconnect the network if it is removed.

Other centrality measures can also be used, such as clustering coefficient, closeness and hop walk statistics, which differ in what type of influence is to be emphasized (Newman, 2010).

### Integration of form and function

The extracted centrality feature vectors can be then combined with function vector (i.e., gene expression) to create a form-function feature matrix 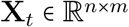, where *n* is the size of Hi-C matrix, *m* is the number of extracted features, and *t* is the time step.

### Data representation on low-dimensional non-linear manifolds

Information redundancy exists in the data matrix 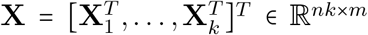, where *k* is the length of time horizon (*k* = 12 in our dataset). For example, the degree centrality and the eigenvector centrality could be correlated, and the replicates of RNA-seq data are strongly correlated. Therefore, data points given by rows of **X** are lying on a manifold with a smaller intrinsic dimensionality *m*′ (often *m′* ≪ *m*) that is embedded in the *m*-dimensional feature space. The goal of dimensionality reduction is to transform dataset **X** into **Y** with lower dimensionality *m′*, while retaining the geometry of the data as much as possible (Van Der Maaten et al., 2009).

Laplacian eigenmap is a non-linear dimensionality reduction technique to find a lowdimensional data representation by preserving local properties of the underlying manifold. We remark that the linear dimensionality reduction technique, principal component analysis (PCA), is also applicable but it cannot adequately handle the nonlinearity embedded in the dataset. The method of Laplacian eigenmaps contain the following steps

- Normalize dataset 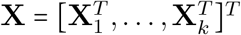 to make different features comparable

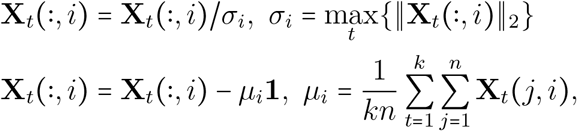

where **X***_t_*(:, *i*) denotes the *i*th column of **X***_t_*, the first transformation ensures that different features are all treated on the same scale, and the second transformation is to zero out the mean of the data.
- Construct a neighborhood graph in which every node is linked with its *p* nearest neighbors. The edge weight is computed using the heat kernel function, leading to a sparse adjacency matrix **W** with entries

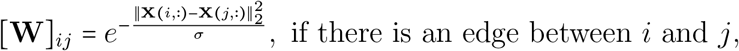

where *σ* is the heat kernel parameter, and we choose *σ* = 200 in our analysis (Van Der Maaten et al., 2009).
- Compute the graph Laplacian matrix **L** = **D** − **W**, where **D** = diag(**W1**). We then solve the generalized eigenvalue problem

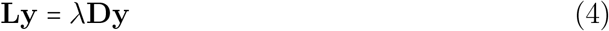

for *m′* smallest nonzero eigenvalues. The resulting eigenvectors 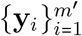 form the lowdimensional data representation **Y** = [**y**_1_, …, **y***_m′_*].

After dimensionality reduction, we can also evaluate the significance of each feature that contributes to the low-dimensional data representation **Y**. Let us consider a linear approximation 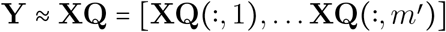, and 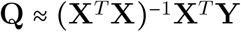. It is clear that there exists a one-to-one correspondence between the column of **Y** and the column of **Q**,

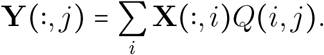

Here *Q*(*i*, *j*) signifies the contribution of the *i*th feature in **X** to the *j*th component of the obtained low-dimensional column-space **Y**. The feature score (FS) for the ith feature corresponding to the *j*th dimension of the subspace is

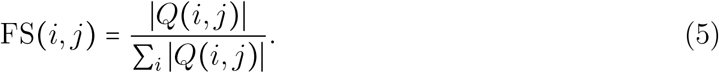

### Fitting the data: minimum volume ellipsoid

The minimum volume ellipsoid (MVE) estimator is the first high-breakdown robust estimator of multivariate location and scatter (Van Aelst and Rousseeuw, 2009). Geometrically, the MVE estimator finds the minimum volume ellipsoid covering, or enclosing a given set of data points. Let 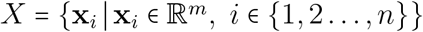 denote the dataset of our interest, where *n* is the number of data points, and *m* is the number of features (or the dimension of the intrinsic low-dimensional manifolds). The ellipsoid that fits into *X* can be parametrized as

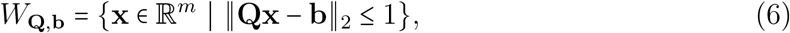

where 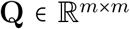 and 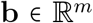 are unknown parameters. The center and the shape of the ellipsoid *E*_**Q,b**_ is given by **c** ≔ **Q**^−1^**b**, and Λ ≔ **Q**^2^ since the ellipsoid (6) can be reformulated as 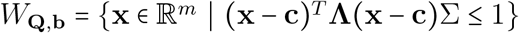. Finding the minimum volume ellipsoid can be then cast as a convex program

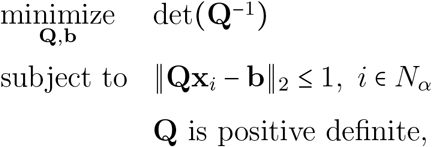

where *N_α_* denotes the set of data within a *α* confidence region, determined by Mahalanobis distances of data below *α* = 97.5% quantile of the chi-square distribution with *l* degrees of freedom (Van Aelst and Rousseeuw, 2009). The MVE estimates the shape of the uncertainty ellipsoid for *X*, which is different from its sample covariance. The latter is the maximum likelihood estimate under the assumption of Gaussian distribution.

### Temporal difference score (TDS)

TDS is introduced to evaluate the temporal difference of form-function characteristics. Let 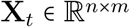 denote data matrix associated with *n* nodes of a network and *m* features. TDS of node *i* at time *t* is defined as

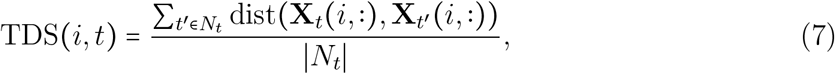

where *N_t_* defines the time window around *t*, namely, *N_t_* = {*t* − 1, *t*}, and dist(·) is a generic distance function between the *i*th row of **X***_t_* and **X***_t′_*. In our analysis, **X***_t_* can represent either network centrality features from Hi-C data or gene expression.

### A/B compartment switching analysis

A/B compartments were identified through methods conceptually similar to that described in (Lieberman-Aiden et al., 2009). Intra-chromosomal Hi-C matrices **H** were binned at the 100-kb level, with unmappable regions and/or regions with no identified contacts removed. Matrices were Toeplitz normalized based on linear genome distance to derive 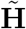 (See Scale-adaptive HiC matrix). The entrywise sign of the principal component of the spatial correlation matrix associated with 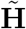 is used to identify A/B compartments.

### Bifurcation: branching trajectory and statistical significance

To depict the branching trajectory at bifurcation, we studied the dataset of human fibroblast proliferation (Chen et al., 2015) and the dataset of the direct cellular reprogramming over a 56-hr time course. First, we found an intrinsic low-dimensional (3D) manifold of centrality-based form-function features under the setting of both proliferation and reprogramming. This is given by the principal subspace of form-function data at the first two time points (corresponding to the fibroblast-like stage). Second, we obtained the 3D data representation of form-function features after projection onto the common subspace for proliferation and reprogramming, and tracked the centroids of fitted ellipsoids (given by MVE estimates) over time. The trajectory of centroids is then smoothed using the cubic spline. Lastly, we provided a statistical significance of the bifurcation, where *P* value is defined from the multivariate Hotelling’s T-Square test associated with the null hypothesis that the centroids of the proliferation and reprogramming are identical at a given time point.

### Bifurcation identification at single gene level

Raw contacts from Hi-C within a ±5 kb window around a gene location are extracted. A {*d* + 1, *d* + 1, *t*} tensor **A***_i,j,t_* is contructed based on the number of MboI cut-sites (GATC) found, *d*, within the region of interest, for each time point sampled *t*. Each element *i*, *j*, *t* of **A** represents the number of contacts found between cut sites {*i* − 1, *i*} and {*j* − 1, *j*} at time t, divided by the total number of contacts found for each time point (RPM). The elementwise difference between time points is calculated, and the summation of the absolute value difference between *t* and *t* + 1 is recorded.

### Identification of genes of interest

Genes of interest (GOIs) are mainly extracted through Gene Ontology (GO), with a few GOI subsets curated through other means. GO-extracted lists include myotube, myoblast, skeletal muscle, fibroblast, and circadian. “Muscle” genes are the union of myoblast, myotube, and skeletal muscle genes. Additional circadian related subsets were extracted from JTK analysis and literature reviews (core circadian), and additional cell cycle subsets were extracted from literature reviews (Table S2).

### Statistical significance of TDS of genes

Give a set of genes, the significance test is made by comparing the average TDS of those genes with a random background distribution. The background distribution is generated by the average TDS of randomly selected same number of genes over 1000 trials. The probability of the right-tailed event is used as *P* value.

### Identification of MYOD/MYOG mediated oscillatory gene expression

Kallisto was used in RNA-seq quantification to obtain TPM (transcripts per million) expression results (Bray et al., 2016). BioCycle was used to identify oscillating transcripts after the identified bifurcation point (32 Hours) with a P value of 0.1 (Agostinelli et al., 2016). Transcripts found to be non-oscillatory before the bifurcation point were identified with a reported P value greater than 0.4. Phase, predicted through a neural network in BioCycle, was used to identify synchronous oscillating transcripts. Synchronous is defined as oscillating transcripts that are in-phase or antiphase within +/− 2 hours. MYOD1 and MYOG gene targets were found by identifying transcription factor binding sites for the respective motifs 10kb upstream or 1kb downstream of transcription start sites (TSS) using MotifMap with a Bayesian Branch Length Score > 1.0 and an FDR < 0.25 (Daily et al., 2011; Xie et al., 2009).

### Super enhancer-promoter region dynamics

SE-P regions for skeletal muscles were downloaded from (Hnisz et al., 2013) (BI_Skeletal_Muscle). The Hi-C contacts between the SE and the associated gene TSS (±1kb) were extracted over time. SE-P contacts were normalized by dividing by the total number of contacts per sample, then multiplying by 100,000,000 (arbitrary scalar to best show trends). To determine the top upregulated genes, the linear regression slope of log2(FPKM) over time was calculated and sorted for each gene.

## DATA AND SOFTWARE AVAILABILITY

The dataset and codes will be reported when the paper is accepted.

## SUPPLEMENTAL ITEM TITLES AND LEGENDS

**Table S1**

Title: Identified genes at A/B switched loci. Related to Figure 3 and S3.

**Table S2**

Title: Gene modules of interest. Related to Figure S4, 4 and 6.

**Table S3**

Title: Gene clusters with significant function and form change during time. Related to Figure 4.

**Table S4**

Title: Core myogenic genes that steer cellular reprogramming. Related to Figure S6.

**Table S5**

Title: JTK output for E-box circadian genes. Related to Figure 6B2.

**Table S6**

Title: List of miRNAs that significantly change expression level over the reprogramming time course. Related to Figure S9.

